# Mrj an Hsp40 family chaperone regulates the oligomerization of Orb2 and long-term memory

**DOI:** 10.1101/2022.10.16.512122

**Authors:** Meghal Desai, Hemant, Ankita Deo, Jagyanseni Naik, Tania Bose, Amitabha Majumdar

## Abstract

Orb2 the Drosophila homolog of Cytoplasmic polyadenylation element binding protein (CPEB) forms prion-like oligomers. These oligomers consist of Orb2A and Orb2B isoforms and their formation are dependent on the oligomerization of the Orb2A isoform. Drosophila with a mutation diminishing Orb2A’s prion-like oligomerization forms long-term memory but fails to maintain it over time. Since, this prion-like oligomerization of Orb2A plays a crucial role in the maintenance of memory, here we aim to find what regulates this oligomerization. In an immunoprecipitation-based screen, we identify interactors of Orb2A in the Hsp40 and Hsp70 families of proteins. Amongst these, we find an Hsp40 family protein Mrj as a regulator of the conversion of Orb2A to its prion-like form. Mrj interacts with Hsp70 proteins and acts as a chaperone by interfering with the aggregation of pathogenic Huntingtin. Unlike its mammalian homolog, we find Drosophila Mrj is neither an essential gene nor causes any gross neurodevelopmental defect. We observe a loss of Mrj results in a reduction in Orb2 oligomers. Further, the knockdown of Mrj in the mushroom body neurons results in a deficit in long-term memory. Our work implicates a chaperone Mrj in mechanisms of memory regulation through controlling the oligomerization of Orb2A and its association with the translating polysomes.

## Introduction

Memory is the experience-dependent ability to preserve and recover information from the past. The molecular mechanism behind long-term memory is long-term potentiation (LTP). LTP is the persistent change observed in synaptic strength accompanied by structural changes like new synaptic growth and stabilization, caused due to repeated patterns of electrical stimulation (Bliss and Lomo, 1973; Harris, 2020; Nicoll, 2017). Several studies ranging across species suggest that protein synthesis plays a crucial role in regulating long-term memory and LTP (Agranoff, 1967; Agranoff et al., 1967; Castellucci et al., 1989; Deadwyler et al., 1987; Eichenbaum et al., 1976; Flexner et al., 1962, 1967, 1965, 1964; Krug et al., 1984; Martin et al., 1997; Stanton and Sarvey, 1984). The translation regulator in Aplysia, Cytoplasmic Polyadenylation Element Binding protein (CPEB) is crucial for the maintenance phase of long-term facilitation and stabilization of learning-induced new synaptic growth (Miniaci et al., 2008; Si et al., 2003a). Its Drosophila homolog Orb2 is necessary for the persistence of long-term memory and has amongst its mRNA targets, genes regulating protein turnover, synapse formation, and neuronal growth (Keleman et al., 2007; Mastushita-Sakai et al., 2010). The mouse homologs CPEB1, CPEB2, and CPEB3 are also implicated in the regulation of memory processes (Berger-Sweeney et al., 2006; Chao et al., 2013; Fioriti et al., 2015; Lu et al., 2017; Stephan et al., 2015).

Aplysia CPEB behaves like functional prions (Heinrich and Lindquist, 2011; Si et al., 2010, 2003b). Prions were discovered as protein-based infectious particles associated with neurodegenerative diseases like Creutzfeldt-Jakob disease, Scrapie, and Bovine spongiform encephalopathy (Prusiner, 2001). Yeast also has several proteins classified as prions (Liebman and Chernoff, 2012; Uptain and Lindquist, 2002; Wickner, 2016; Wickner et al., 1999, 1996). Prion proteins exist in two distinct conformational and functional variants: one monomeric and another oligomeric amyloid-like form. The oligomeric form is dominant and self-perpetuating as it can convert the monomeric to the amyloid form. Due to Aplysia CPEB’s prion-like characteristics, it is suggested that synaptic stimulation causes CPEB to convert to its prion-like state and this state can self-sustain as long as monomers are getting synthesized. The prion-like oligomers can further regulate the protein synthesis of the target mRNAs needed for the maintenance of long-term memory. This model gets support from studies with Drosophila Orb2, where a point mutation in Orb2 that disrupted its prion-like oligomerization caused an impairment in the persistence of long-term memory (Hervás et al., 2016; Majumdar et al., 2012). Biochemically separated monomeric and oligomeric forms of Orb2 exhibit functional differences from each other. In *in vitro* translation assays with its target mRNAs, the monomer was found to act as a translational repressor by decreasing its poly-A tail length while the oligomer acted as a translation activator by elongating its poly-A tail (Khan et al., 2015). This observation is similar to the idea that prions based on their conformational difference and oligomeric status can have different biochemical functions. These translation-enhancing oligomers when visualized using cryo-electron microscopy, appear as amyloids and like prions can seed the conversion of the monomers to the amyloid form (Hervas et al., 2020)

Orb2 has two isoforms, Orb2A and Orb2B, both containing the common prion-like domain and a C terminal RNA binding domain. The two isoforms differ in the position of the prion-like domain, enrichment in the brain, and propensity to aggregate. Orb2A has 8 amino acids while Orb2B has 162 amino acids, in front of the prion-like domain. In comparison to Orb2B, Orb2A is less abundant in the brain. Orb2A has a higher propensity to aggregate compared to Orb2B. Though both the Orb2A and Orb2B isoforms interact amongst themselves and are present in the oligomers, it is Orb2A that acts as a seed for Orb2 oligomerization. This is supported by several lines of evidence. Firstly, a deletion of Orb2A causes a drastic reduction in the formation of endogenous Orb2 oligomers in the brain. Secondly, a point mutation of the fifth Phenylalanine to Tyrosine (F5Y) in Orb2A reduces its ability to oligomerize and interferes with the maintenance of long-term memory (Majumdar et al., 2012). This residue is specific to only Orb2A and is not present in Orb2B. Thirdly, genetic deletion experiments suggested Orb2A does not need its RNA binding domain and Orb2B does not need its prion-like domain for the maintenance of memory (Krüttner et al., 2012). Finally, the addition of the prion-like domain of Orb2A can seed monomeric Orb2B to oligomerize and result in its change to a translational activator (Khan et al., 2015). All this evidence suggests oligomerization of Orb2A is crucial for the formation of Orb2 oligomers and the maintenance of memory. Hence in this study, we asked, what regulates the oligomerization of Orb2A, how this regulator affects the overall Orb2 oligomers in the brain, and if this regulator plays any role in long-term memory.

Taking a cue from the Yeast prion literature, where the protein folding machinery/ chaperones act as key regulators of prions, we hypothesize, chaperones may also play a role in the regulation of Orb2A oligomerization. In Yeast, the Hsp70, Hsp40, and Hsp104 chaperones have been found to regulate the oligomerization and propagation of prions (Chernova et al., 2017; Liebman and Chernoff, 2012; Summers et al., 2009; Tuite and Lindquist, 1996; Wickner, 2016). Using an immunoprecipitation-based screen and a yeast-based prion conversion assay, here we identify Drosophila Mrj as a regulator of Orb2A’s prion-like conversion. Mrj stands for the mammalian relative of DnaJ (Izawa et al., 2000) and functions as a chaperone in mammals. We find Drosophila Mrj to behave similarly to mammalian Mrj as a chaperone and interfere with the aggregation of pathogenic Huntingtin. While knockout of Mrj in mice is embryonic lethal (Hunter et al., 1999), in Drosophila we find it not to be an essential gene and observe this knockout to have a reduced amount of Orb2 oligomers. We further find Mrj is needed in specific mushroom body neurons for long-term memory and mechanistically it may have a role in regulating Orb2A’s association with the polysomes.

## Results

### Identification of chaperone interactors of Orb2

We started with identifying homologs of the Yeast Hsp70, Hsp40, and Hsp104 families of proteins. The Drosophila genome lacks any Hsp104 homolog but it has members of the Hsp40 and Hsp70 classes of proteins (Ayme and Tissières, 1985; Raut et al., 2017; Velazquez et al., 1980). So, for our screen to identify Orb2A regulators, we decided to focus on these two groups. The Hsp40 group of proteins is classified as proteins containing DnaJ domains. The Drosophila genome contains 39 such genes (Supplementary Figure 1). For the Hsp70 class, Drosophila has both the heat shock inducible Hsp70 (Hsp68, Hsp70Aa, Hsp70Ab, Hsp70Ba, Hsp70Bb, Hsp70Bc, and Hsp70Bbb) and the constitutively expressing Hsc70 (Hsc70-1, Hsc70-2, Hsc70-3, Hsc70-4, Hsc70-5, and Hsc70Cb) (Supplementary Figure 2). We made a library of 37 of the Hsp40 genes tagged with HA epitope and 4 of the Hsp70 genes tagged with Flag epitope. The 4 Hsp70 genes were selected based on their sequence variability from each other. We next transfected each of these constructs with Orb2A in Drosophila S2 cells and used these cells to perform immunoprecipitation with an anti-Orb2 antibody (Figure 1A, C). The immunoprecipitate was probed in a western blot for the presence of HA-tagged Hsp40 or Flag-tagged Hsp70 protein. From this immunoprecipitation-based screen, we observed the proteins CG4164, CG9828, DroJ2, CG7130, Tpr2, Mrj, Hsc70-1, Hsc70-4, Hsc70Cb, and Hsp70Aa as interactors of Orb2A (Figure 1B, D). For 31 Hsp40 proteins, we could not detect interaction with Orb2A in our immunoprecipitation screen (Supplemental Figure 3).

**Figure 1:**
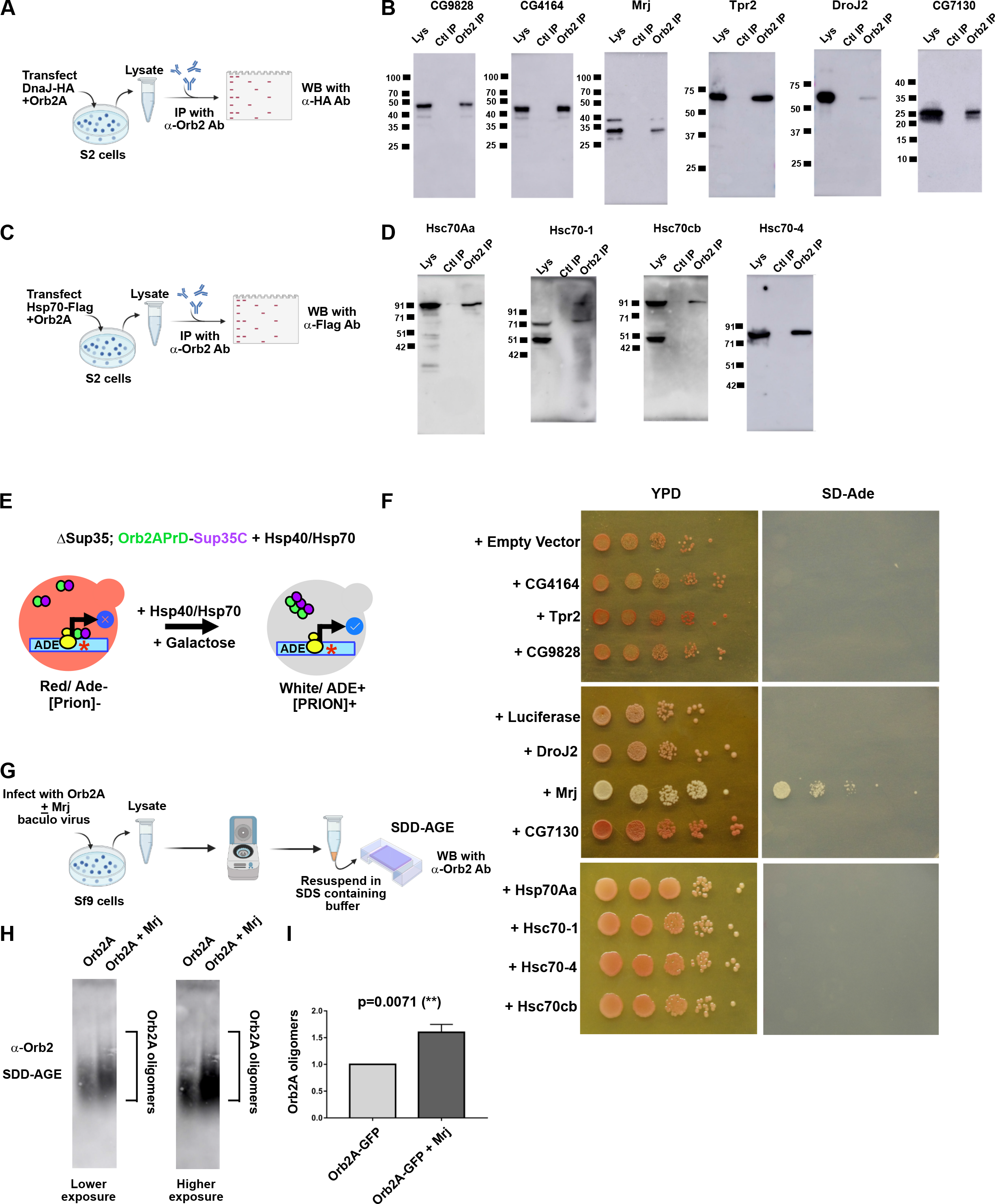
Identification of Mrj as a regulator of Orb2A’s prion-like oligomerization. **A**. Schematic of immunoprecipitation assay to identify Hsp40 interactors of Orb2A. Individual constructs coding for members of the Hsp40 family with HA tag were cotransfected in S2 cells with Orb2A construct and immunoprecipitation was performed with anti Orb2 antibody. The immunoprecipitate was probed with anti HA antibody to detect if the Hsp40 proteins were interacting with Orb2A. **B**. From 37 proteins of the Hsp40 family, the immunoprecipitation (IP) screen identified CG4164, CG9828, Tpr2, DroJ2, Mrj, and CG7130 as interactors. Lys is the input lysate, Ctl IP is the control IP with beads only and Orb2 IP is the IP with anti-Orb2 antibody **C**. Schematic of immunoprecipitation assay to identify Hsp70 interactors of Orb2A. Individual constructs coding for members of the Hsp70 family with Flag tag were cotransfected in S2 cells with Orb2A construct and immunoprecipitation was performed with anti Orb2 antibody. The immunoprecipitate was probed with anti Flag antibody to detect if the Hsp70 proteins were interacting with Orb2A **D**. All four proteins of the Hsp70 family screened in the immunoprecipitation experiment, Hsp70Aa, Hsc70-1, Hsc70Cb, and Hsc70-4 were found to be interacting with Orb2A **E**. Schematic of the Yeast based screen. A Sup35 knockout strain rescued by a chimeric construct expressing Orb2A’s prion-like domain tagged with the C-terminal domain of Sup35 is transformed with galactose inducible Hsp40 and Hsp70 constructs. A premature stop codon in the Ade 1-4 gene is used as a reporter. When the Orb2PrD-Sup35C protein is in non-prion form, it helps the ribosomes encountering the premature stop codon in Ade1-4 to fall off leading to no read-through translation. The colonies for this non-prion state are red and cannot grow in Adenine deficient media. If the Orb2APrD-Sup35C protein gets converted to a prion-like state, it will fail to dislodge the ribosomes encountering the premature stop codons, and as a result, read-through translation will happen and now the cells will be white, which can now grow in adenine deficient media. The screen consists of inducing the chaperones with galactose in the Prion negative strain and screening for their ability to change the color to white and grow in adenine-deficient media **F**. Galactose induction of individual chaperones of both Hsp40 and Hsp70 family of proteins in Prion negative Orb2APrD-Sup35C strain, caused only Mrj to convert the Prion negative state of the cells to Prion positive state, as evidenced by the change in the colony color to white in YPD media and causing it to grow now in adenine **G**. Schematic of SDD-AGE assay to quantitate the change in Orb2A-GFP oligomerization in presence of Mrj. Sf9 cells were infected with viruses for Orb2A alone and Orb2A with Mrj. The lysate from these cells was centrifuged and the resulting pellet was resuspended in an SDS containing buffer, subjected to SDD-AGE, and further probed with anti-Orb2 antibody **H**. Representative SDD-AGE blots showing increased levels of Orb2A oligomers in presence of Mrj **I**. Quantitation of Orb2A oligomers in presence and absence of Mrj. Data is represented as a relative fold change for Orb2A in presence of Mrj as compared to without Mrj. Data is represented as Mean ± SEM and significance is tested using unpaired t*-*test.

### Drosophila Mrj converts the prion-like domain of Orb2A from non-prion to a prion-like state

We next asked if any or all of these interactors can regulate the oligomerization of Orb2. Toward this, we tested these genes in a heterologous yeast chimeric Sup35-based system. Sup35 is a translation terminator which can exist in both non-prion and prion forms (Cox, 1965; Liebman and Chernoff, 2012; Paushkin et al., 1997; Stansfield et al., 1995). Replacement of Sup35’s prion-like NM domain with putative prion-like domains of other proteins was previously used to identify several new prions (Alberti et al., 2009; Halfmann et al., 2011). In this assay when the NM domain is replaced with the N terminal 162 amino acids of Orb2A (Orb2A-PrD), the prion-like behavior could be visualized as a red (non-prion) or white (prion) colony color. While the white colonies grow in Adenine deficient media, the red colonies are unable to grow in the same (Hervás et al., 2016). To check what effect the Orb2 interactors will have on the non-prion state of Orb2A, we made Galactose inducible constructs for expressing the ten Orb2 interactors and transformed these into red-colored, Adenine negative, Orb2A-PrD-C-Sup35 strain. The transformed colonies were independently grown and then induced with Galactose (Figure 1E) and plated with serial dilutions on complete media (YPD) and Adenine deficient media. Out of all the ten genes from the Hsp40 and Hsp70 groups and controls, we observed only Mrj coexpression could change the red color of the colony into white. These cells could also now grow in Adenine deficient media (Figure 1F). This yeast-based screen of interactors of Orb2 suggested Mrj could convert the non-prion form of Orb2A to its prion-like state.

A conversion of Orb2A from its non-prion to prion-like form suggests an increase in its oligomeric state. We tested this in Sf9 cells by coexpressing Orb2A with and without Mrj. The cells were lysed and centrifuged and the pellet was further resuspended in an SDS-containing buffer and then resolved on a semi-denaturing detergent agarose gel electrophoresis (SDD-AGE) (Figure 1G) (Alberti et al., 2009; Halfmann and Lindquist, 2008). SDD-AGE has been previously reported to detect SDS-resistant Orb2 oligomers as a smear in agarose gels (Khan et al., 2015; L. Li et al., 2016; Majumdar et al., 2012). On probing the SDD-AGE blot with an Orb2 antibody we observed a significant increase in Orb2 oligomers in presence of Mrj (Figure 1H, I). This suggests that Drosophila Mrj can increase the oligomerization of Orb2A.

### Drosophila Mrj like its mammalian homolog interacts with Hsp70 proteins and can act as a chaperone preventing aggregation of pathogenic Huntingtin

The amino acid sequence of Drosophila Mrj is well conserved with its homologs in humans, mouse, rat, frog, and zebrafish (Supplemental Figure 4A). We asked if Drosophila Mrj has similar properties to Mammalian Mrj. Similar to mammalian Mrj (Izawa et al., 2000), Drosophila Mrj is also present in both the cytoplasm and nucleus (Supplemental Figure 4B). Mammalian Mrj is reported to self-interact to form oligomers (Karamanos et al., 2019; Månsson et al., 2014; Söderberg et al., 2018). We tested the self-association of Drosophila Mrj in vivo by coexpressing Mrj-RFP and Mrj-HA in S2 cells and immunoprecipitating Mrj-RFP (Figure 2A). On probing the immunoprecipitate, we could detect the presence of Mrj-HA in it (Figure 2B), suggesting that Drosophila Mrj self-associates inside cells. We also observed purified recombinant Mrj precipitate within a few days of purification suggesting the formation of higher-order oligomers that eventually phase out of the solution. Using Dynamic light scattering of purified recombinant Mrj, we noticed different sized Mrj populations in solution, which over three days moved to higher size suggesting the self-association of the protein (Supplemental Figure 4C).

**Figure 2:**
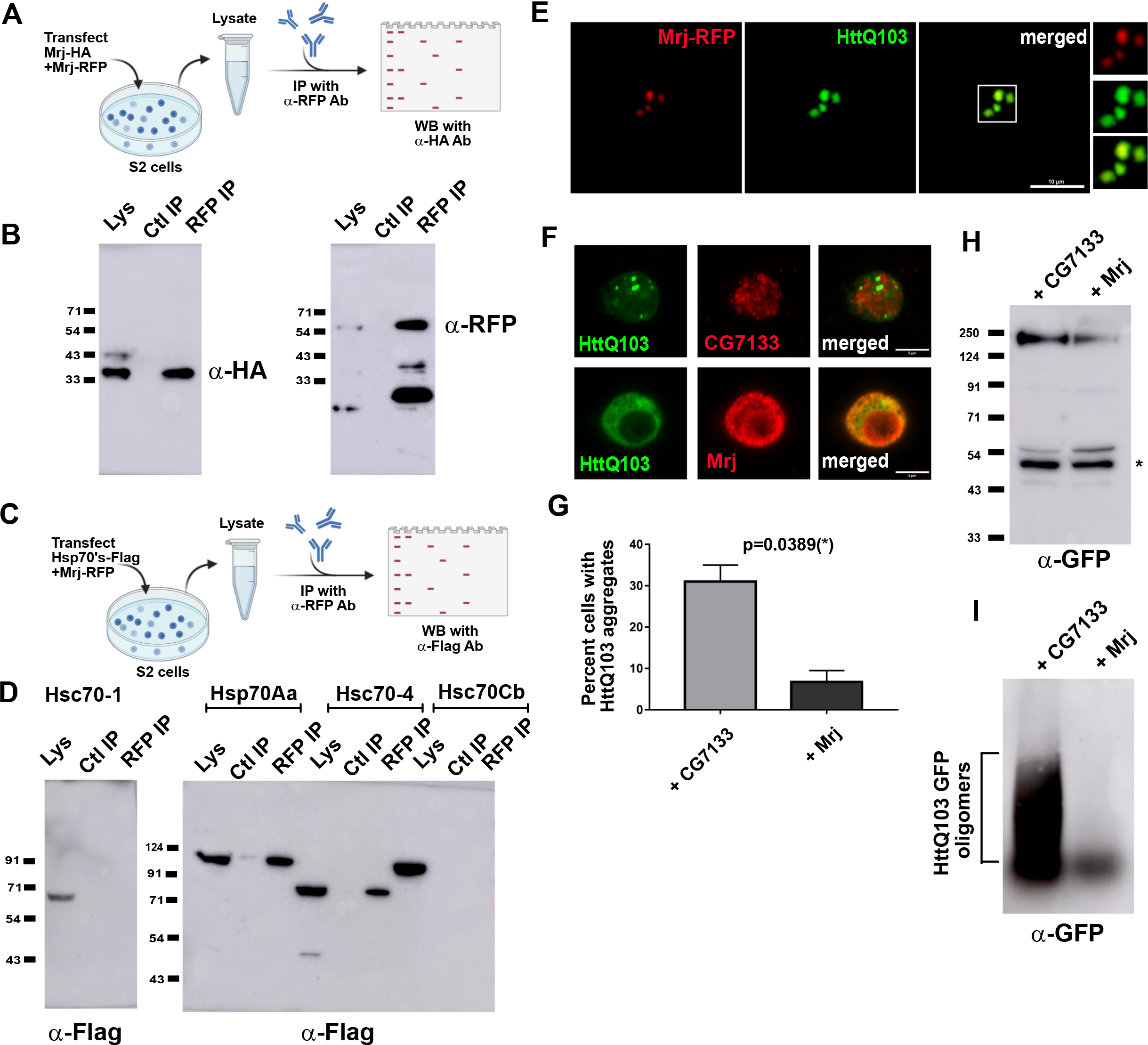
Drosophila Mrj behaves like its mammalian homolog as a chaperone preventing Huntingtin aggregation. **A**. Schematic of immunoprecipitation assay to check the possibility of self-oligomerization of Mrj. Two Mrj constructs, Mrj-HA and Mrj-RFP were cotransfected in S2 cells. The cells were lysed and immunoprecipitation was done with anti-RFP (RFPTrap beads). The immunoprecipitate was next probed with anti-HA antibody **B**. Left panel shows a western blot of immunoprecipitated Mrj-RFP probed with anti-HA antibody showed the presence of Mrj-HA suggesting interaction between Mrj-RFP and Mrj-HA. The right panel shows the same blot probed with anti-RFP antibody to confirm the pulldown of Mrj-RFP. **C**. Schematic of immunoprecipitation assay to identify Hsp70 interactors of Mrj. Flag-tagged Hsp70 constructs were cotransfected with Mrj-RFP. Lysate from these cells was used for immunoprecipitation with anti-RFP beads, and the immunoprecipitate was probed with anti-Flag antibody **D**. In IP experiment, of the 4 Hsp70’s tested here, Hsp70 Aa and Hsc70-4 show their presence in the immunoprecipitate suggesting of their interaction with Mrj **E**. Confocal image of S2 cell coexpressing Mrj-RFP and Httexon1Q103-GFP shows colocalization of Mrj with Htt aggregates **F**. Representative images of Httexon1Q103-GFP cells coexpressing with CG7133-HA and Mrj-HA suggest a decrease in the Htt aggregates in presence of Mrj. The Mrj-HA constructs are more efficient in decreasing Htt aggregation compared to Mrj-RFP **G**. Quantitation of the percentage of HttQ103exon1GFP expressing cells with aggregates in presence of CG7133 and Mrj suggests a significant decrease of Htt aggregates in presence of Mrj. Data is represented as Mean ± SEM **H**. Western blot of lysates from S2 cells coexpressing Httexon1Q103-GFP with Mrj and CG7133 shows similar amounts of Htt (monomer size marked with *) in SDS-PAGE **I**. SDD-AGE from S2 cell lysate coexpressing Httexon1Q103-GFP along with CG7133 and Mrj showed a decreased amount of Htt oligomers in presence of Mrj.

The Hsp40 chaperones interact with the Hsp70 family of proteins and act as cofactors for different substrates (Kampinga and Craig, 2010). Mammalian Mrj is reported to interact with Hsp70 proteins (Izawa et al., 2000). Co-immunoprecipitation experiment of Drosophila Mrj with four of the Drosophila Hsp70 proteins suggested Drosophila Mrj can interact with at least two of these proteins Hsp70Aa and Hsc70-4 (Figure 2C, D).

Mammalian Mrj could inhibit the aggregation and toxicity of Huntingtin Exon1 protein containing 150 Q repeats (Chuang et al., 2002). Drosophila Mrj was found to colocalize with expanded poly Glutamine aggregates in the brain and could rescue its toxicity in the Drosophila eye (Fayazi et al., 2006). We tested the effect of Drosophila Mrj on Htt oligomers and aggregates using biochemical and imaging-based assays. Towards this, we coexpressed a Htt construct spanning the exon1 with 103 Q repeats (Zhang et al., 2010) and observed colocalization of Mrj-RFP with HttQ103 aggregates in S2 cells (Figure 2E). On coexpression of Mrj-HA with HttQ103-GFP, we observed a significant decrease in the number of cells showing visible aggregates in comparison to coexpression with a control DnaJ domain-containing protein CG7133 (Figure 2F, G). While there was no detectable difference in total Htt levels in the lysates of these two sets in SDS-PAGE (Figure 2H), in SDD-AGE, we observed a clear reduction of Htt oligomers in presence of Mrj (Figure 2I). We also had a similar observation of Mrj decreasing Htt aggregate and oligomers (Supplementary Figure 4D, E, F), for a longer Htt construct spanning the caspase cleaved 588 amino acid fragment (Joag et al., 2020; Weiss et al., 2012)). Together the observations of Drosophila Mrj interacting with Hsp70 proteins and its ability to interfere with pathogenic Htt aggregate formation suggests its possible role as a chaperone.

### Drosophila Mrj knockout is viable and does not show any gross developmental defect

We next used the CRISPR-Cas9 system to generate a knockout of Drosophila Mrj by introducing a Gal4 cassette in the Mrj locus (Figure 3A). We confirmed the knockout by using both PCR and western blotting with an anti-Mrj antibody (Figures 3B and C). The mammalian Mrj knockout mice were reported to be lethal at the mid-gestational stage at embryonic 8.5 days due to failure in chorioallontoic fusion (Hunter et al., 1999). In the knockout, the Keratin filaments had undergone collapse due to the formation of Keratin inclusion bodies which disrupted the chorionic trophoblasts leading to cellular toxicity (Watson et al., 2007). Apart from the Keratin filament disorganization Mrj knockouts also exhibited defects in Actin cytoskeleton organization, misexpression of E-cadherin and β catenin, and disorganization of Extracellular matrix (ECM) (Watson et al., 2011). Here, we found the Drosophila Mrj knockout line to be homozygous viable. Developmentally there was no defect in any organization of any body structure, and there was no sterility associated with the homozygous line. In terms of overall cellular organization, we observed no defect in the actin cytoskeleton and the gross organization of the mushroom body as evidenced by staining with Phalloidin and anti-FasII antibody (Figure 3D). We next checked if the absence of Mrj caused endogenous proteins to aggregate in the brain. Towards this, we looked for the Drosophila homolog of p62, Ref(2)P which is a regulator of protein aggregates and is present in ubiquitinated protein aggregates associated with aging and neurodegenerative disorders (Nezis et al., 2008). On Ref(2)P immunostaining to label ubiquitinated protein aggregates, we could not detect any difference in the labelling between the wild type and Mrj knockout brains (Figure 3E). We also stained the wild type and Mrj knockout brains with ubiquitin antibody, and again saw no difference in ubiquitin labelling in the brains (Figure 3F).

**Figure 3:**
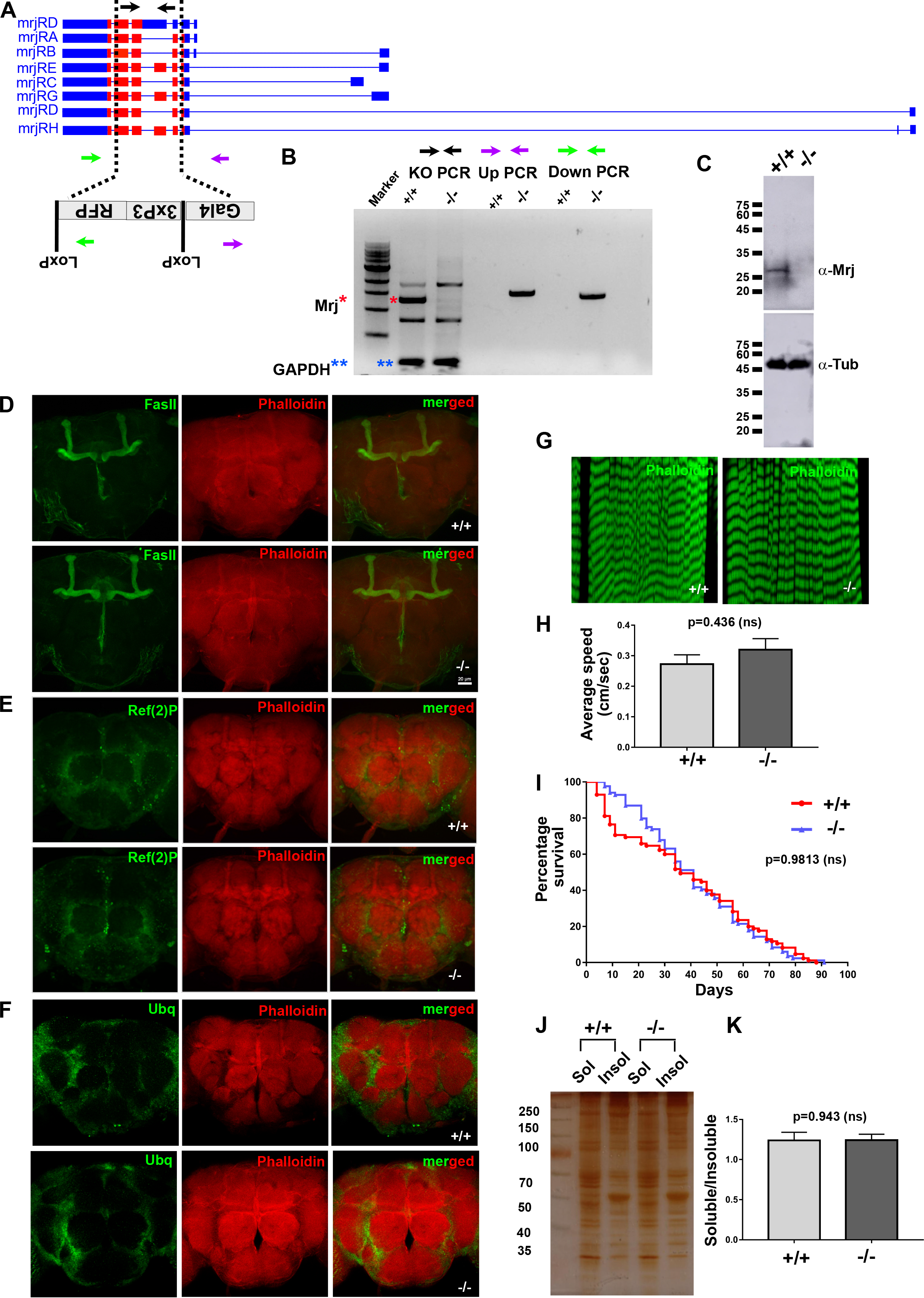
Drosophila Mrj is not an essential gene unlike its mammalian homolog. **A**. Schematic of the genomic organization of the Mrj gene and the knockin of the Gal4-loxP-3XP3-RFP-loxP cassette in the locus to make Mrj knockout. The black arrows represent the PCR primers for confirming the Mrj knockout, and the green and purple arrows represent the PCR primers for up and down PCR to check the knockin of the cassette in the Mrj locus **B**. Confirmation of Mrj knockout using genomic DNA PCR with the knockout, up and down PCR primers. The red * marks the amplified band for Mrj in the wild type (+/+)and its absence in the Mrj knockout (−/−). GAPDH PCR amplification (blue **) was used as the loading control in the PCR **C**. Confirmation of Mrj knockout using western blot using an anti-Mrj antibody. Anti-α-Tubulin antibody was used as the loading control **D**. Immunostaining of wild type (+/+) and Mrj knockout (−/−) with anti-FasII antibody and Phalloidin shows no gross difference in the overall morphology of mushroom body and the brain **E**. Immunostaining of wild type (+/+) and Mrj knockout (−/−) with anti-Ref(2)P antibody and Phalloidin and **F**. with anti-Ubiquitin antibody and Phalloidin shows no gross difference between the two sets **G**. Phalloidin staining of muscles from third instar larvae of wild type (+/+) and Mrj knockout (−/−) shows no gross difference **H**. Quantitation of the average speed of wild type (+/+) and Mrj knockout (−/−) shows no significant difference between the two. Data is represented as Mean ± SEM and unpaired t test is used to check significance **I**. Kaplan-Meier survivor curve shows no significant difference in the life span of wild type (Red) and Mrj KO (Blue) **J**. Representative image of silver-stained gel of soluble and insoluble protein fractions, from wild type (+/+) and Mrj knockout (−/−) fly heads **K**. Quantitation of soluble to insoluble protein ratio from wild type (+/+) and Mrj knockout (−/−) flies showed no significant difference. Data is represented as Mean ± SEM and Data is represented as Mean ± SEM and unpaired t test is used to check significance.

As mutations in human Mrj are associated with Limb-Girdle muscular dystrophy (LGMD), we checked if the muscles in Mrj knockout fly show any defect in terms of organization or degeneration. On staining the larval muscles with phalloidin, we observed no evidence of degeneration or disorganization in the Mrj knockout compared to the wild type (Figure 3G). We next checked the adult Mrj knockout fly for any defect in locomotion and found them to show no significant difference in their walking speed in comparison to the wild type (Figure 3H). This ruled out the possibility of any drastic muscle defect later in development. In terms of life span, we observed no difference between the knockout and wild-type animals (Figure 3I). Since Mrj is known to prevent the formation of pathogenic Htt aggregates, we next asked if Mrj knockout animals show any difference in the total protein distribution in the brain between the soluble and insoluble fractions in comparison to the wild type. We lysed fly heads in Triton X-100 containing lysis buffer and separated the supernatant as the soluble fraction. The remaining pellet was solubilized using an SDS containing lysis buffer and separated as an insoluble fraction. We resolved these fractions in SDS-PAGE, performed silver staining (Figure 3J), and quantitated the ratio of soluble/ insoluble protein. Here also we observed no significant difference in terms of the amount of soluble/insoluble protein ratio between the wild type and Mrj knockout animals (Figure 3K). These results together suggested that under normal growing conditions, Mrj knockout animals were indistinguishable from wild-type ones in terms of their development, brain and muscle organization, and locomotion abilities. Also, for ubiquitinated aggregates and insoluble protein amounts no difference could be seen between the two sets.

### Drosophila Mrj interacts with Orb2A independent of its RNA binding domain

On co-transfecting Mrj-RFP with Orb2A-GFP we could observe colocalization between Orb2A punctae and Mrj in 27 percent of cotransfected cells (Figure 4A). To address which region of Orb2A is needed for its interaction and colocalization with Mrj we cotransfected different deletion mutants of Orb2A with Mrj-RFP and performed imaging and co-immunoprecipitation (Figure 4D) experiments. Upon co-transfecting the Orb2A325-GFP construct which lacks the C terminal RNA binding domain with Mrj RFP, we observed colocalization in 35 percent of cotransfected cells (Figure 4B). This suggested the interaction between Mrj and Orb2A is independent of its RNA binding domain. This is further supported by the observation that immunoprecipitation of Mrj from these cells could pull down Orb2A325 (Figure 4E). We also cotransfected Mrj with a Δ162Orb2A-GFP construct which lacks the prion-like domain. On imaging we could not detect any colocalization (Figure 4C). and in immunoprecipitation experiments, observed the inability of Mrj to pull down Δ162Orb2A (Figure 4F). This suggested the interaction of Mrj with Orb2A to be dependent on the N terminal 162 amino acid long region of Orb2A containing the prion-like domain. This also supported the previous observation of Mrj converting the Orb2A-PrD-Sup35 from a non-prion to a prion-like state, where the N terminal 162 amino acids of Orb2A were used to make the chimera with Sup35.

**Figure 4:**
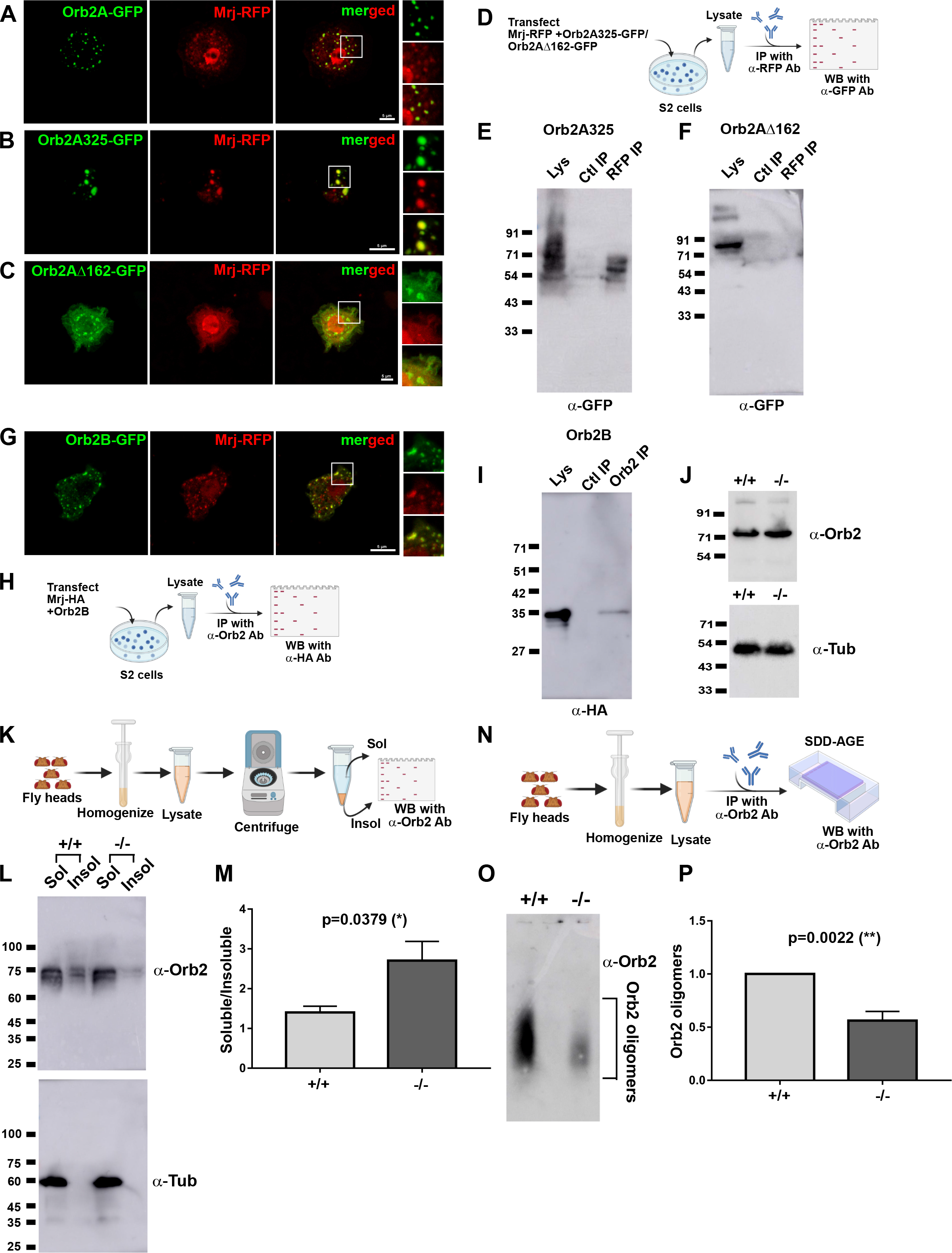
Mrj knockout shows reduced Orb2 oligomers. Representative image of S2 cells co-expressing Orb2A-GFP and Mrj-RFP showing colocalization of Mrj with Orb2A aggregates **B**. Representative image of S2 cells co-expressing Orb2A325-GFP (RNA binding domain deleted construct) and Mrj-RFP showing colocalization of Mrj with Orb2A 325 aggregates. **C**. Representative image of S2 cells co-expressing Orb2AΔ162-GFP (prion-like domain deleted construct) and Mrj-RFP showing no colocalization of Mrj with Orb2AΔ162 **D**. Schematic of immunoprecipitation assay to check for interaction between Mrj-RFP and Orb2A325-GFP and Orb2AΔ162-GFP **E**. Immunoprecipitatated Mrj-RFP pulls down Orb2A325 suggesting the interaction between Mrj and Orb2A is independent of its RNA binding domain **F**. No interaction detected between Orb2AΔ162-GFP and Mrj-RFP in immunoprecipitation experiment suggesting the interaction of Orb2A with Mrj is dependent on the prion-like domain of Orb2A **G** Representative image of S2 cells co-expressing Orb2B-GFP and Mrj-RFP showing colocalization of Mrj with Orb2B aggregates **H**. Schematic of immunoprecipitation assay to check for interaction between Mrj-HA and Orb2B-GFP **I**. Immunoprecipitation experiment showed interaction between Orb2B and Mrj **J**. Western blot of fly head extracts from wild type and Mrj KO shows similar amounts of Orb2B in both of them. α-Tubulin is used as a loading control here **K**. Schematic of the process of preparing the soluble and insoluble fractions from fly head extract to check for differential distribution of Orb2B **L**. Western blot of soluble and insoluble fractions from wild type and Mrj KO fly heads show a reduced amount of Orb2B in the insoluble fraction of Mrj knockout. α-Tubulin was used as the loading control **M**. Quantitation of relative Orb2B level ratio in soluble to insoluble fractions show a significant increase in Mrj KO flies as compared to control wild type flies. Data is represented as Mean ± SEM. **N**. Schematic of the Orb2 pulldown experiment followed by SDD-AGE to check for Orb2 oligomers **O**. Representative SDD-AGE blot shows reduced amount of Orb2 oligomers in Mrj knockout **P**. Quantitation of Orb2 oligomers in Mrj knockout and wild type. Data is represented as Mean ± SEM and unpaired t-test is used to check significance.

### Drosophila Mrj interacts with Orb2B and its knockout shows reduced amounts of Orb2 oligomers

We also looked at the other Orb2 isoform Orb2B. Though Orb2B-GFP expressing cells rarely show punctate appearance characteristic of aggregation, these punctae showed colocalization with Mrj-RFP (Figure 4G). In immunoprecipitation experiments, on pulling down Orb2B, we detected Mrj in the immunoprecipitate (Figure 4H, I). Overall, these experiments suggest Mrj interacts with both the Orb2A and Orb2B isoforms, and this interaction is independent of the RNA binding domain.

As the endogenous Orb2 oligomers in the Drosophila brain consist of both Orb2A and Orb2B, we asked what happens to endogenous Orb2 oligomers in Mrj knockout animals. In western blots from fly heads, the detectable form is Orb2B representing its relatively higher abundance in the brain. We could not detect any significant difference in Orb2B levels between the wild type and Mrj knockout (Figure 4J). Next, we separated the soluble and insoluble fractions from wild-type and Mrj knockout fly heads and probed them with an anti-Orb2 antibody. In this assay, while we could detect the presence of Orb2B in both the soluble and insoluble fractions in the wild-type fly head lysate, in the Mrj knockout, Orb2B was more enriched in the soluble fraction (Figure 4K, L, M). This suggested lesser amounts of Orb2B in the insoluble fraction in Mrj knockout indicating lighter or lesser Orb2 oligomers. We further tested this by immunoprecipitating Orb2 from wild-type and Mrj knockout animals and subjecting the immunoprecipitate to SDD-AGE (Figure 4N). On probing the SDD-AGE blots, we observed the presence of significantly reduced levels of Orb2 oligomers in Mrj knockout brains in comparison to the wild type (Figure 4O, P). Based on these results we interfere that the chaperone Mrj is probably regulating the oligomerization of endogenous Orb2 in the brain.

### Mrj is needed in the mushroom body for the regulation of long-term memory

As our observations imply Orb2 oligomerization is dependent on Mrj, we next asked if Mrj is expressed in brain structures that are relevant for memory. Towards this, we used the Mrj knockout line, where we introduced a Gal4 cassette in the locus. On using this line to express CD8GFP, we observed labeling in the mushroom body lobes (Figure 5A). In the Drosophila brain, the mushroom body is the center for the formation and storage of memory.

**Figure 5:**
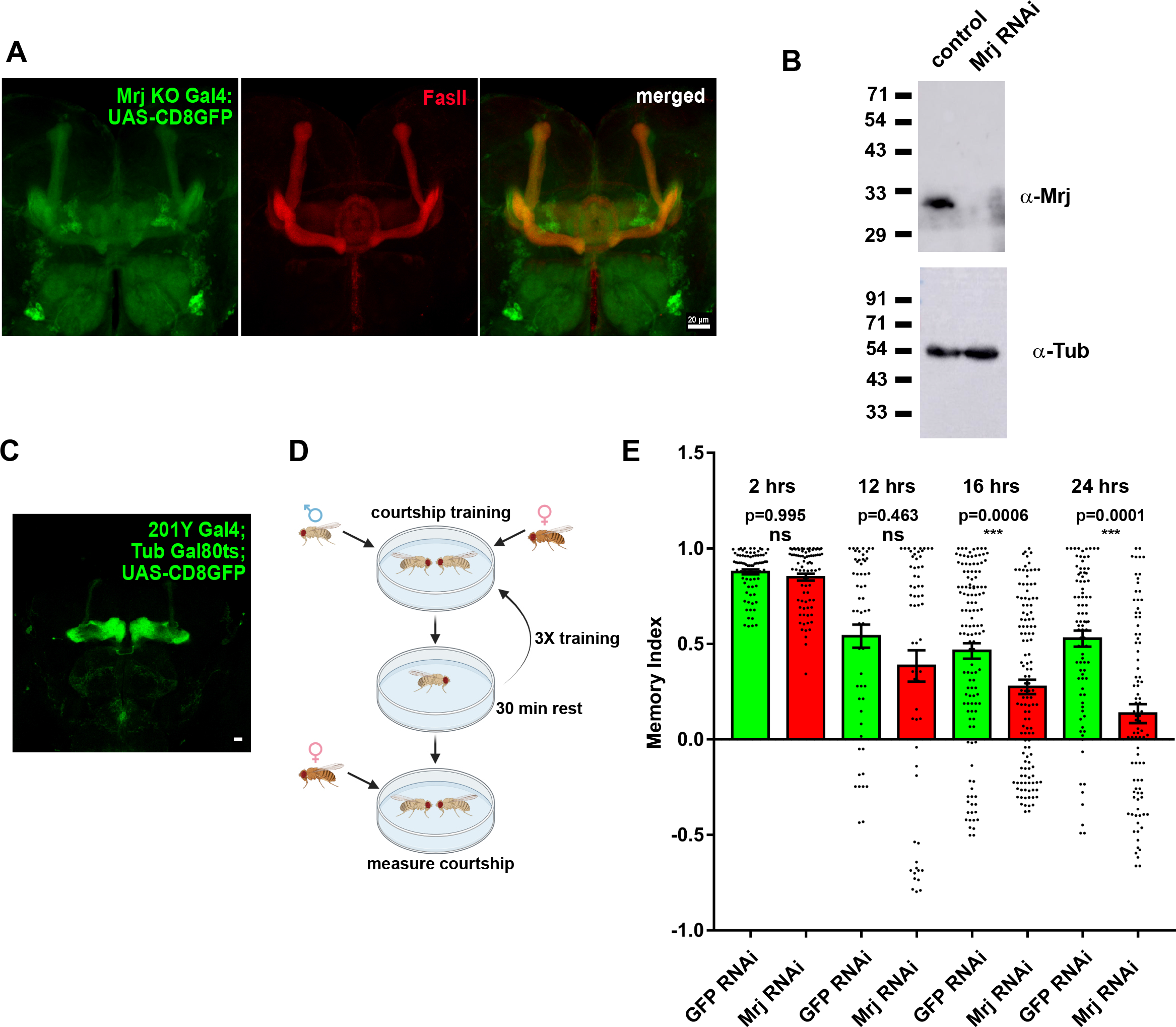
Knockdown of Mrj in mushroom body neurons causes a long-term memory deficit. **A**. Representative image of **the** expression pattern of Mrj KO Gal4 visualized using the reporter CD8GFP (green) and counterstained with anti-FasII antibody (Red) shows the expression of GFP in mushroom body neurons **B**. Confirmation of knockdown of Mrj by pan-neuronal expression of an Mrj RNAi line using a western blot with anti-Mrj antibody. Anti-α-Tubulin antibody was used as the loading control **C**. Representative image of the expression pattern of 201Y Gal4 visualized using the reporter CD8GFP **D**. Schematic of Male courtship suppression-based memory assay **E**. Knockdown of Mrj in specific mushroom body neurons using 201Y Gal4 causes a significant memory deficit in comparison to the control from 16 hours onwards. Data are represented as Mean ± SEM and Mann-Whitney test is done to test for significance.

To understand the role of Mrj in the mushroom body, we used an Mrj RNAi line. We first checked for the ability of the Mrj RNAi line to knock down Mrj in the brain. On inducing Mrj RNAi using a pan-neuronal Elav Gal4 driver, we could not detect Mrj in the RNAi fly head extract in comparison to the control (Figure 5B). This indicated the effective knockdown of endogenous Mrj to non-detectable levels.

Using this RNAi line we now asked, if Mrj is specifically required in the mushroom body neurons for memory. We used a mushroom body-specific 201Y Gal4 line which extensively expresses in the γ lobes along with some neurons of the α/β lobes (Figure 5C) to knock down Mrj in mushroom body neurons. Previously, the expression of Orb2 transgenes using this 201Y Gal4 line was found to be sufficient to rescue memory deficits in Orb2 mutants (Keleman et al., 2007). Hence this Gal4 line provides us with the advantage of knocking down Mrj only in the Orb2 relevant neurons. Though earlier we could not detect any gross developmental defects in Mrj knockout flies, we wondered if there might be fine undetected developmental defects. To rule out memory deficiency caused due to such defects, we decided to perform the Mrj knockdown in the adult stage after all development has taken place. Towards this, we coupled the 201Y Gal4 line with a temperature-sensitive Gal80 ts (McGuire et al., 2004, 2003) and used this to perform the Mrj knockdown with the inducible Mrj RNAi line. These flies were allowed to lay eggs, grow and eclose at 22° C temperature. At this temperature, the Gal80 would suppress the transcriptional activity of Gal4 and the Mrj RNAi would not be induced. Post eclosion, these flies were shifted to 30° C, where the Gal80 becomes inactive resulting in restoration of the transcriptional activity of Gal4 causing the Mrj RNAi to knockdown Mrj in the mushroom body neurons. We next subjected these mushroom body-specific Mrj knockdown animals to courtship suppression-based memory assay (Figure 5D). Male flies when subjected to repeated rejections by mated females, suppress their courtship behavior post-training. Compared to wild-type animals, mutants defective in their ability to retain memory show a faster recovery of the courtship behavior suggesting a decrease in their capacity to remember. At post-training time points of 2 hours and 12 hours, we could not detect any significant difference in memory scores between the control and Mrj knockdown groups, suggesting a normal formation of early and intermediate-term memory. However, at the 16-hour time point, the memory score became significantly reduced for the Mrj knockdown animals in comparison to the control animals (Figure 5E). This difference became even greater at the 24-hour time point where the memory score for Mrj knockdown decreased drastically compared to the control (Figure 5E). Together these experiments suggest Mrj is needed in mushroom body neurons for regulating long-term memory in Drosophila.

### Mrj regulates the association of Orb2A with translating ribosomes

Since long-term memory is dependent on protein synthesis, and the knockdown of Mrj in mushroom body neurons causes a long-term memory deficit, we asked if Mrj plays any role in this protein synthesis. We first addressed if Mrj can associate with ribosomes. Hence Mrj was cotransfected with Rpl18-Flag, which was earlier found to be incorporated in assembled ribosomes and can be used to pulldown polysomes (Khan et al., 2015). On immunoprecipitating Rpl18 with anti-Flag antibody, we could detect Mrj in the immunoprecipitate (Figure 6A) suggesting Mrj most likely interacts with the translating ribosomes. We next performed polysome fractionation of Mrj expressing S2 cell extract in a 5-40% sucrose gradient. On probing the fractions, we found Mrj to be present in the polysome fractions (Figure 6B). However, on doing the fractionation for EDTA treated extract, which disassembles the polysomes, we observed Mrj is still present in all the fractions but now getting more enriched and shifting to the heavier fractions (Figure 6C). One possible explanation for this is, Mrj forms oligomers which are probably of similar density as the polysomes. As this doesn’t explain the shift of Mrj to heavier fractions on EDTA treatment, an alternate possibility is that the polysomes associated with Mrj, stabilize it and this prevents Mrj from forming heavier oligomers. On EDTA treatment this association is disrupted and free Mrj gets destabilized and forms higher-sized oligomers through increased self-association.

**Figure 6:**
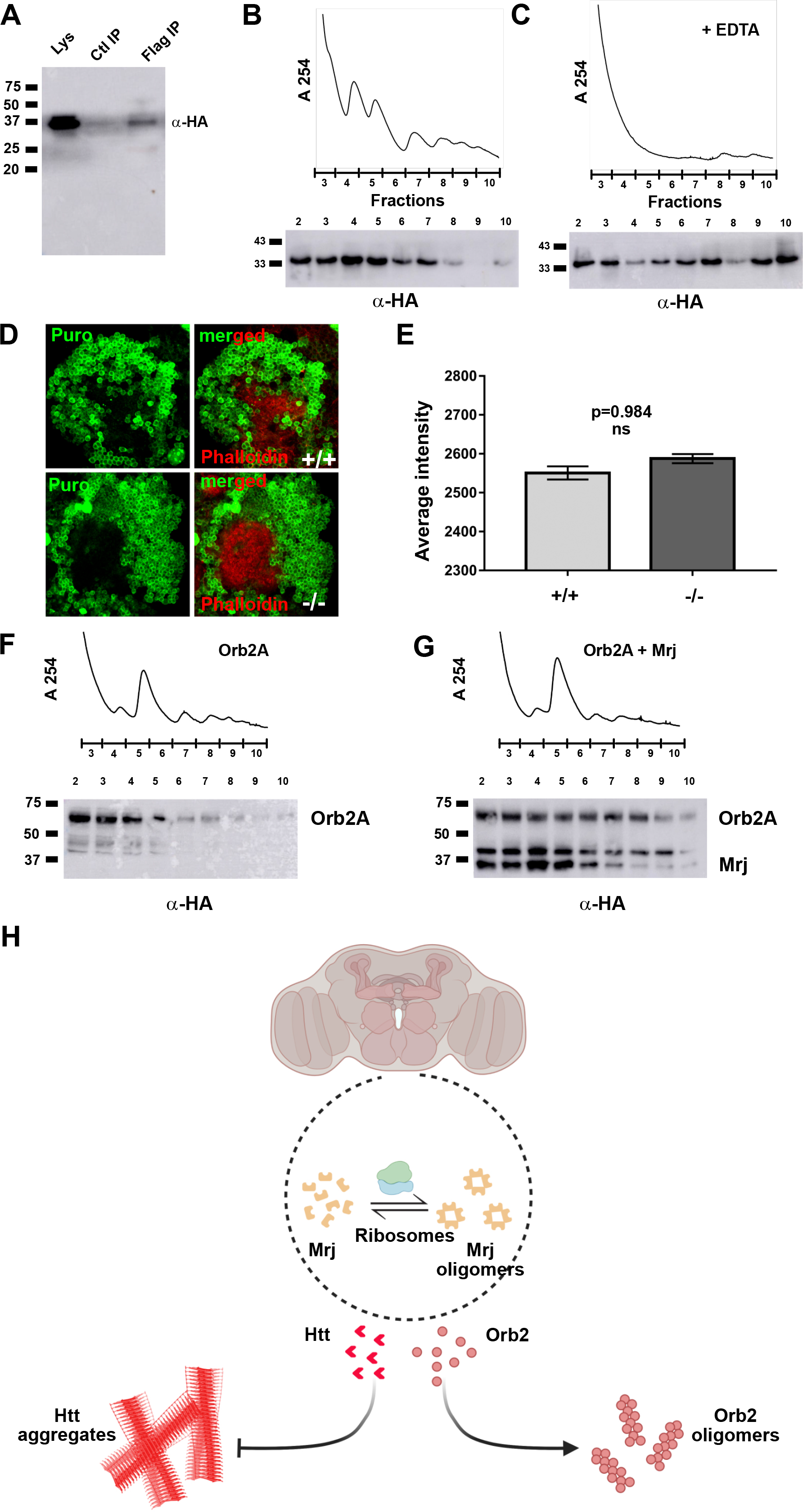
Mrj interacts with ribosomes and helps in regulating Orb2A’s association with polysomes. **A**. Immunoprecipitated Rpl18-Flag which is incorporated in ribosomes pulls down Mrj, suggesting Mrj is associated with ribosomes **B**. Upper panel shows a representative polysome profile of lysate obtained from cells expressing Mrj-HA. The numbers in X-axis represent the fraction number. The lower panel is a western blot of individual fractions, showing the presence of Mrj in polysome fractions **C**. Upper panel shows a representative polysome profile of EDTA treated lysate obtained from cells expressing Mrj-HA. The numbers in X-axis represent the fraction number. The lower panel is a western blot of individual fractions, showing the shift of Mrj to heavier fractions in comparison to non-EDTA treated lysate **D**. Representative images of mushroom body Kenyon cells immunostained with anti-Puromycin antibody to detect newly translated proteins. Red shows the counterstain with Phalloidin **E**. Quantitation of anti-Puromycin staining intensity shows no significant difference between the wild type (+/+) and Mrj knockout (−/−) **F**. Upper panel shows representative polysome profile of lysate obtained from cells expressing Orb2A-HA. The numbers in X-axis represent the fraction number. The lower panel shows a western blot detecting Orb2A in individual fractions **G**. Upper panel shows a representative polysome profile of lysate obtained from cells co-expressing Orb2A-HA and Mrj-HA. The numbers in X-axis represent the fraction number. The lower panel shows a western blot detecting Orb2A and Mrj in individual fractions. In comparison to only Orb2A, in presence of Mrj, more Orb2A is detected in polysome fractions **H**. Schematic of the model originating from this work. We hypothesize association of Mrj with Ribosomes is probably stabilizing it and regulating its oligomerization. Mrj acts as an inhibitor of aggregation of pathogenic Huntingtin (Htt) and positively regulates the formation of Orb2 oligomers.

As Mrj is associated with the polysomes, we next checked if Mrj played any role in global protein synthesis. For this, we used the Puromycin incorporation assay to compare protein synthesis between wild-type and Mrj knockout brains(Deliu et al., 2017; Schmidt et al., 2009). Puromycin gets incorporated into translating proteins due to its similarity with tRNA and blocks further incorporation of amino acids. The amount of Puromycin incorporation can be visualized using immunostaining with an anti-Puromycin antibody. Using this assay, we did not observe any significant difference in Puromycin incorporation between the brains of wild type and Mrj knockout (Figure 6D, E). This suggests the absence of Mrj does not cause any gross defect in general translation.

We next addressed the effect of Mrj on the association of Orb2A with the translating ribosomes. We transfected S2 cells with Orb2A along with and without Mrj and used these cells to perform polysome fractionation. On assaying the different fractions with Orb2 antibody, we observed that in presence of Mrj, more Orb2A is present in the polysome fractions (Figure 6F, G). This indicates that in presence of Mrj, the association of Orb2A with the translating ribosomes is enhanced.

## Discussion

One key mechanism involved in the regulation of the persistence of memory is the prion-like activity of CPEB in Aplysia (Heinrich and Lindquist, 2011; Miniaci et al., 2008; Raveendra et al., 2013; Si et al., 2010, 2003a, 2003b) and Orb2 in Drosophila (Keleman et al., 2007; Khan et al., 2015; Krüttner et al., 2015, 2012; Majumdar et al., 2012). In this work, we identify Mrj as a protein that is regulating the conversion of Orb2A from non-prion to prion-like oligomeric form. The mammalian homolog of Mrj was earlier reported to be acting as an oligomeric chaperone interfering with the aggregation of Huntingtin aggregates (Chuang et al., 2002; Hageman et al., 2010; Kakkar et al., 2016; Rodríguez-González et al., 2020; Thiruvalluvan et al., 2020). Mrj was also observed to be upregulated in the astrocytes of Parkinson’s patients and helps in suppressing α-Synuclein aggregation, suggesting an overall protective mechanism against aberrant aggregation of proteins causing neurodegeneration (Aprile et al., 2017; Durrenberger et al., 2009). Like its mammalian homolog, we found Drosophila Mrj to be an oligomeric protein interacting with Hsp70 chaperones, which can decrease Huntingtin aggregates in cells. This along with previous reports of Drosophila Mrj rescuing effects of polyglutamine aggregates (Bason et al., 2019; Fayazi et al., 2006) suggests that Drosophila Mrj acts similarly to its mammalian homolog as a chaperone.

Knockout of Mrj in mice caused embryonic lethality, suggesting Mrj to be an essential gene. Mammalian Mrj was previously implicated in LGMD where patients with missense mutations in the gene showed the presence of rimmed vacuoles and inclusion bodies in the muscle (Bengoechea et al., 2015; Harms et al., 2012; Sarparanta et al., 2012). In contrast to mammalian Mrj-associated phenotypes, we found Drosophila Mrj knockout to be homozygous viable and having no signs of defects in the development of the brain, cytoskeleton, and muscle. There was no difference with the wild-type control in terms of life span and accumulation of Ref(2)P and Ubiquitinated proteins, which are markers of aging and aggregates in the brain. While the mice knockout for Mrj is embryonic lethal what makes the Drosophila Mrj knockout to be so different from the observed phenotype of mammalian Mrj knockout? Mammalian Mrj was found to be an interactor of Keratin protein and mouse Mrj knockout exhibited Keratin aggregates which might have contributed to the lethality. In Drosophila, since there is no Keratin homolog present, we speculate that this probably helps the Mrj knockout to escape lethality.

We find reduced Mrj knockout to have reduced Orb2 oligomers in comparison to wild-type suggesting a requirement of Mrj in regulating the formation of endogenous Orb2 oligomers. Our findings indicate that Mrj’s behavior towards Orb2 is very different from its known protective, anti-aggregating function against pathogenic proteins associated with neurodegenerative diseases. While for both Huntingtin and Synuclein, deletion of Mrj caused an increase in their aggregation, in the case of Orb2, deletion of Mrj decreased its aggregation. This suggests the same chaperone may have different effects on the regulation of aggregation for pathogenic against non-pathogenic functional amyloids.

How does the Mrj-dependent regulation of the prion-like nature of Orb2, compares with other studies of the effect of Hsp40 chaperones with prions and RNA-binding proteins? Studies in yeast showed the maintenance and propagation of three prions associated with Rnq1, Ure2, and Sup35 are dependent on an Hsp40 family protein Sis1 (Aron et al., 2007; Higurashi et al., 2008; Lopez et al., 2003). Sis1 acts as a disaggregase in collaboration with Hsp104 and Hsp70 by helping in the fragmentation of the oligomers of these prions to generate propagating seeds (Nakagawa et al., 2022). Another Hsp40 family protein Ydj1 together with Hsp70 has been found to inhibit the formation of Ure2 prion state in yeast by delaying its aggregation to form amyloid fibrils (Lian et al., 2007; Sharma et al., 2009). Apart from the regulation of prions, the chaperones Sis1 together with Hsp104 and Hsp70 regulate the dispersal of the heat-induced aggregates of an RNA-binding protein Pab1 (Yoo et al., 2022). Overexpression of mammalian Mrj/DnaJB6 was able to cure Ure2 prions by helping in the dissolution of the aggregates in cells (Reidy et al., 2016; Zhao et al., 2018). This suggests the chaperones including Mrj have an anti-aggregation role for prion aggregates whereas, in the case of Orb2, this is promoting its aggregation. The only Hsp40 chaperone which was found similar to Mrj in increasing Orb2’s oligomerization is the yeast Jjj2 protein.

Our observations of knocking down Mrj in the specific mushroom body neurons suggest Mrj plays a role in these neurons for the regulation of long-term memory. Some previous studies indicated that chaperones might help in the regulation of memory. In the context of neurodegenerative diseases, a DnaJ domain-containing protein Droj2 has been suggested to be a negative regulator of memory in the Drosophila model of Alzheimer’s disease (Ring et al., 2022). ER chaperones Hspa5 and small heat shock protein 22 (sHsp22) were found to improve the impairment in spatial learning and memory deficit in a mice Tauopathy model (Chatterjee et al., 2022; Rodriguez Ospina et al., 2022). Through our work we implicate the chaperone Mrj, to play a crucial role in the regulation of long-term memory in a non-stressed and non-disease condition.

In this study, we also try to find a plausible mechanism of Mrj-dependent memory regulation. As protein synthesis is vital for long-term memory regulation and Orb2 is a translation regulator, we asked if Mrj plays any role in this. We observed Mrj interacts with ribosomes and is present in the polysome fractions. This observation is also supported by proteomics-based interactome analysis of human Mrj/ DnaJb6, where it was found to interact with several ribosomal proteins (Piette et al., 2021). Mammalian Mrj exists in an equilibrium between its monomers and oligomers and interfering with its oligomerization reduces its anti-aggregation property on Htt aggregates (Kakkar et al., 2016; Karamanos et al., 2020). We observe that on disrupting the polysomes, Mrj migrated to heavier fractions. Based on this, we hypothesize the association of Mrj with the polysomes is probably regulating its oligomerization and stabilizing them and preventing it from forming bigger oligomers.

There are also examples from yeast where DnaJ domain-containing Hsp40 family proteins, Zuotin1, Sis1, and Jjj1, and Hsp 70 family proteins Ssa1 and Ssz1 have been reported to associate with ribosomes (Horton et al., 2001; Meyer et al., 2007, p. 1; Yan et al., 1998). Zuo1 and Ssz1 together are referred to as Ribosome associated chaperones (RAC) and together with Ssb, they interact with the newly translated protein coming out of the ribosome tunnel and help in its folding. Since the loss of Mrj does not cause any deficit in global protein synthesis, our findings suggest that Mrj is not a “generalist” but a “specialist” who is most probably involved in regulatory roles for only certain substrates. In this case, it regulates the oligomerization of Orb2A, and polysome assays suggest it probably regulates its association with the translating ribosomes.

While we have identified Mrj as the chaperone regulator of Orb2 oligomerization, Orb2 is probably not the only substrate of Mrj. Another known RNA binding protein interactor of Mrj is Hrb98DE the Drosophila ortholog of human hnRNPA1 and hnRNPA2B1 associated with inherited myopathies (S. Li et al., 2016). There might also be other RNA-binding proteins as the substrate of Mrj. Different DnaJ domain proteins have been reported to interact amongst themselves and form functional disaggregation complexes (Nillegoda et al., 2015). Whether Mrj also interacts with other Hsp40 family proteins and forms functional complexes in the regulation of Orb2 and long-term memory needs to be addressed in the future.

## Methods and materials

### Construction of library to express Hsp40 and Hsp70 proteins in S2 cells and yeast

The Drosophila Hsp40 and Hsp70 proteins were PCR amplified using gene-specific primers and were cloned into a Topo-D-Entr vector using TopoD-Entr cloning kit (Invitrogen). Sequence-confirmed constructs were then transferred to destination vector pUASg-HA (Hsp40 genes) with 3X HA tag in the C terminal (Bischof et al., 2013) and pUASt-ccdB-FLAG (pTWF) with 3X FLAG tag in the C terminal (For Hsp70’s) using LR clonase, and these clones were used for all S2 cell-based experiments. Mrj-RFP construct was made by transferring Topo-D-Entr-Mrj construct to the destination vector pAc5.1-ccdB-RFP (pAWR) using LR clonase. For expression of Hsp40s and Hsp70s in Yeast, Topo-D-Entr clones were cloned in destination vector pAG424-GAL-ccdB-HA with 3X HA tag in the C terminal. Orb2A-Sup35C chimera was prepared by transferring a Topo-D-Entr construct of Orb2A (corresponding to N terminal 162 amino acids) to the destination vector pAG415-ADH1-ccdB-SUP35C using LR clonase.

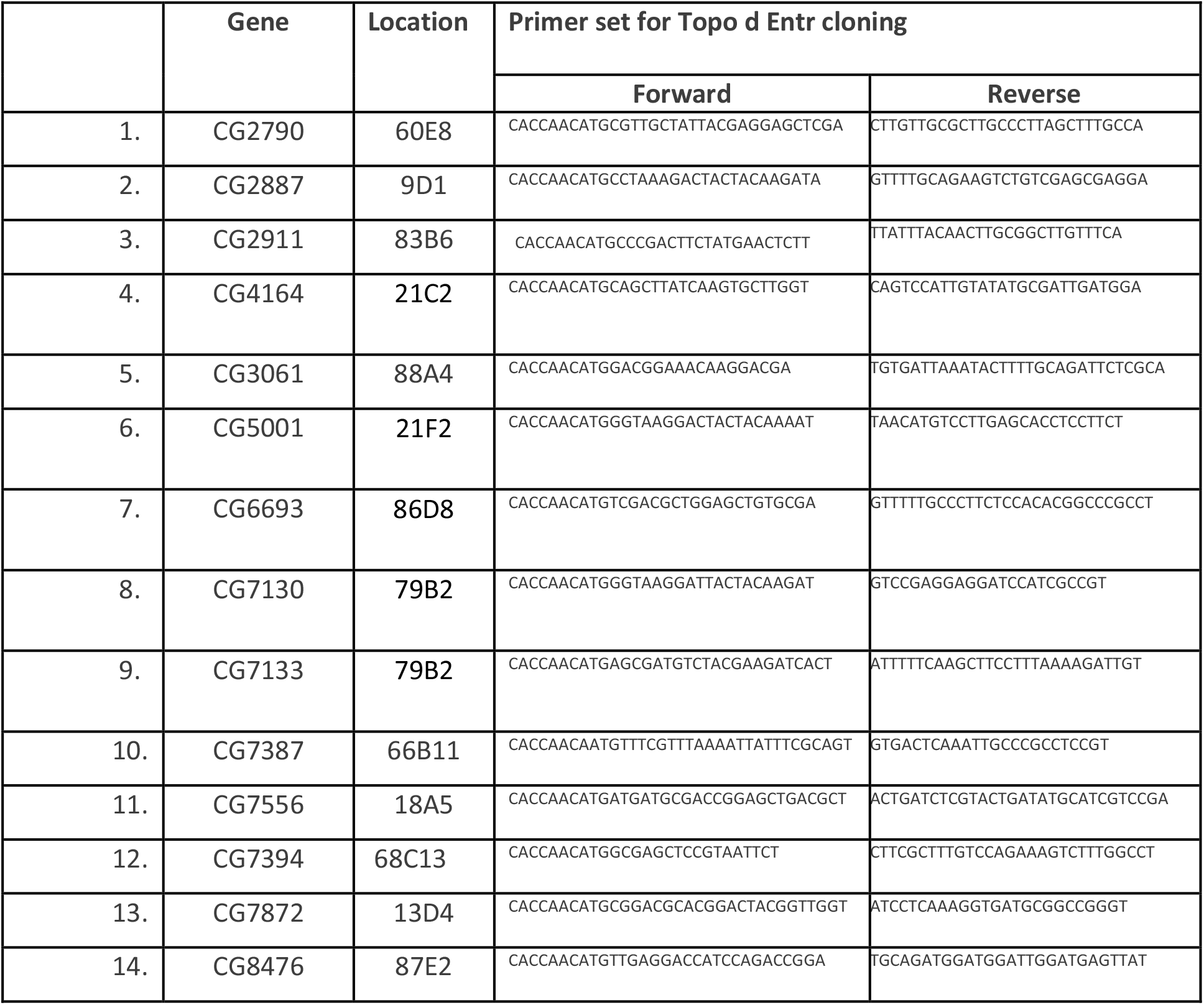
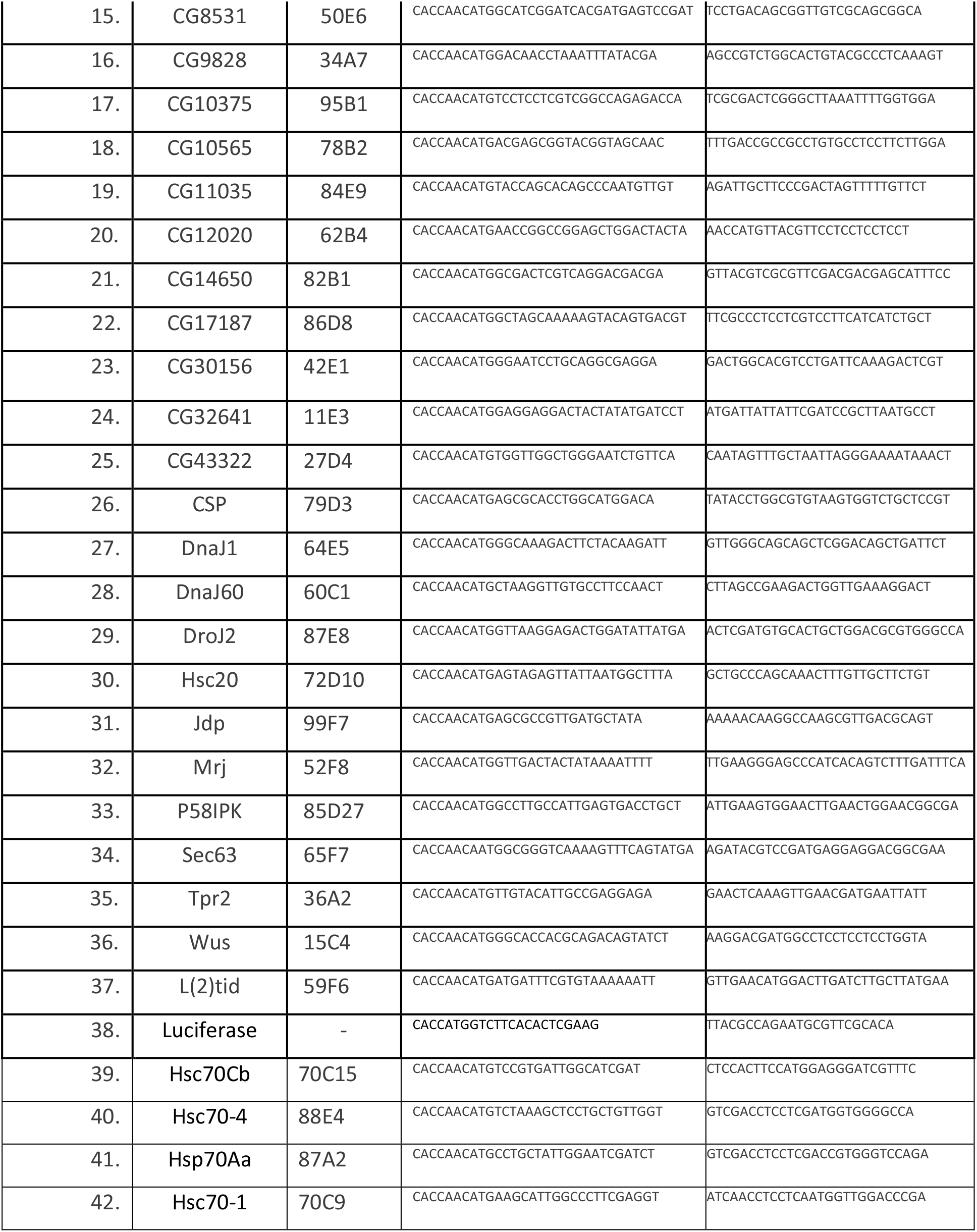

### Construction of truncated Orb2A constructs to express in S2 cells

Truncated Orb2A fragments amplified by the designated primers were cloned in Topo-D-Entr and then transferred to the destination vector pUASt-ccdB-GFP (pTWG) using LR clonase.

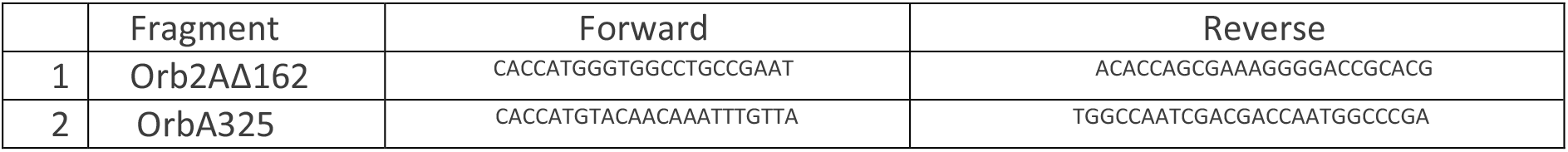

### Construction of Baculo virus for expressing Mrj and Orb2

The Mrj-HA fragment was excised from pUASg-Mrj-HA construct with NotI and NheI digestion and cloned in pFastBac1 digested with the same enzymes. For untagged Orb2A, the Orb2A fragment was PCR amplified and cloned in pFastBac1, for the GFP-His tagged Orb2A, the same was cloned in pFastBac-GFP-His vector. These pFastBac based constructs were next transformed in DH10Bac cells and positive colonies were screened using X-Gal based Blue white screening. While colonies were further grown and Bacmid were purified. These Bacmids were next transfected in Sf9 cells using Cellfectin (Invitrogen) to generate Baculo virus, which were confirmed for expression using western blots.

### S2 cell culture and transfection

S2 cells were grown in Schneider’s media supplemented with 10% FBS. Transfections of plasmids were done with Effectene using the manufacturer’s protocol.

### Immunoprecipitation from S2 cells

Transfected S2 cells were lysed in lysis buffer (150 mM NaCl, 50mM Tris, 1% NP40, and protease inhibitors) were centrifuged at 10000g for 10 minutes at 4°C. The supernatant was separated and incubated with antibodies for 2 hours at 4°C. Immunoprecipitation was performed by binding this lysate with pre-blocked (with 1% BSA) Protein-A beads for an additional 2 hours and further collecting the beads by spinning at 1000g for 2 min. The beads were next washed three times for 10 min with cold lysis buffer. Proteins bound to the beads were eluted by the addition of 0.2 M Glycine (pH 2) for 1 min and further neutralization with 1.5 M Tris (pH 9). The eluates were next mixed with loading dye and processes for western blots. Validation of the Orb2 antibodies is shown in Supplementary Figure 5A, B, C, and D. Validation of the Orb2 antibody in the immunoprecipitation experiments is shown in Supplementary Figure 5E, and F.

### Western blot

The immunoprecipitated samples were boiled for 10 minutes and were directly loaded and ran on polyacrylamide gels (10%, 12%, or 15% gels according to molecular weight of the protein of interest) and further electro-blotted onto a PVDF membrane for 3h at 80V in cold wet transfer buffer. The membranes were blocked with 5% nonfat dry milk in TBST buffer and incubated with respective primary and HRP-conjugated secondary antibodies. The membrane was washed with TBST and incubated with chemiluminescence reagents. The membrane was next put between two transparency sheets and the chemiluminescent signals on it were imaged using a GE Healthcare Lifesciences AI600 Imager.

### Yeast prion assay

A yeast strain bearing a Sup35 knockout rescued by a Sup35 expressing plasmid sup35∷HygB pAG426GPD-SUP35 (Mata, leu2-3,112;his3-11,-15;trp1-1;ura3-1;ade1-14;can1-100;[RNQ+];sup35∷HygB;pAG426GPD-Sup35) was transformed with pAG415 ADH-RA162-Sup35C/Leu plasmid and selected on SD-Leu plates. The colonies were grown in YPD media with repetitive media changes/transfer and then plated/streaked on 5-FOA plates, as Ura marker-containing cells would not grow on this selection media resulting in the selection of colonies without the pAG426GDP-Sup35 rescue plasmid. Colonies growing in 5-FOA plates were further re-streaked in Leu media to check and confirm the presence of pAG415 ADH-RA160-Sup35C/leu plasmid. These colonies were also checked by streaking them in YPDG media to confirm they were not petites. Positive colonies were further streaked and grown on YPD media and checked for colonies of red and white color. Isolated red and white colonies were restreaked in SD-Ade media to confirm only the white colonies but not the red colonies are capable of growing in this. The red colonies which are prion negative were further grown and transformed with pAG424Gal-Hsp40/Hsp70/Trp constructs. As control empty 424 Gal vector and 424-Gal-luciferase vectors were used. Colonies were selected on SD-Ura-Trp plates. Single colonies were grown in Sd-Ura-Trp media overnight. The next day cultures were spun down, given 2 washed with PBS, and resuspended in Raf-Ura-Trp media by adjusting OD600 to 0.2. Cultures were grown to the mid-log phase and induced by the addition of 20% Galactose. After 24 hours of induction, cultures were serially diluted and spotted on YPD and SD-Ade plates.

### Generation of recombinant Mrj

Primers MrjFwd_NdeI (GCGGCAGCCATATGTATGGTTGACTACTATAAAAT) and MrjRev_XhoI (GTGGTGGTGCTCGAGTTGAAGGGAGCCCATCA) were designed with a 15bp overhang from pET28a vector. Mrj was PCR amplified using Q5 polymerase (NEB) using TopoD-Entr-Mrj clone as a template. The amplicon was ligated with pET28a vector digested with Nde1-HF and Xho1-HF enzymes (NEB) using the Infusion cloning kit (Takara). Ligations were transformed in DH5α cells plated on LB+ Kanamycin plates. Sequence confirmed plasmid was transformed in BL21-DE3 cells and grown in LB+Kan media at 37°C under shaking conditions till an OD of 0.6. The culture was next shifted to 30°C and induced with IPTG (final concentration of 0.5 mM) for 6 hours. The cells were next pelleted with centrifugation and lysed in denaturing lysis buffer (8M Urea, 100mM Na_2_PO_4_, 10mM tris-Cl, 10mM imidazole) with sonication. The lysate was centrifuged at 10000g for 10 minutes and the supernatant was allowed to bind with 1 ml of Ni-NTA beads (Qiagen) for 1 hour. Post binding the beads were washed with denaturing wash buffer (8M Urea, 100mM Na_2_PO_4_, 10mM tris-Cl, 40mM imidazole). The wash buffer was changed from denaturing wash buffer to native wash buffer(100mM Na_2_PO_4_, 300mM NaCl, 40mM imidazole) by a gradual decrease of Urea in the wash buffer. Mrj protein bound to these beads was next eluted using native elution buffer (50mM Na_2_PO_4_, 300mM NaCl, 300mM imidazole). This purified protein was used for raising antibodies and downstream DLS experiments.

### Mrj Antibody generation

To generate antibodies, purified Mrj was mixed with Freund’s complete and incomplete Adjuvants and injected in guinea pig over a 72-day immunization protocol followed by the collection of blood and separation of serum. The serum was used in western blots with wild-type and Mrj knockout fly head extracts to confirm the specificity of the Mrj antibody.

### Generation of Mrj Knockout

Upstream gRNA sequence CTTCACTTCACTATCGGTAG[CGG] and downstream gRNA sequence ACTTCGACGGTGTTTGTGAA[TGG] were cloned into U6 promoter plasmid. Cassette Gal4-RFP containing Gal4, SV40 polyA terminator, two loxP sites, 3xP3-RFP, and two homology arms were cloned into pUC57-Kan as donor template for repairing Mrj. The targeting gRNA construct along with hs-Cas9 and the donor plasmid were together microinjected into embryos of control strain w[1118]. F1 flies carrying the selection marker of 3xP3-RFP were further validated by genomic DNA PCR and sequencing. This resulting line is a deletion allele of Mrj, with a knockin of Gal4 cassette in the Mrj locus.

### Soluble-Insoluble fractionation

Fly heads of Mrj-KO and W118 flies were collected and homogenized in lysis buffer (50mM Tris pH7.5, 150mM NaCl, 0.1% Triton-X-100). These lysates were centrifuged at 14,000g for 15mins at 4°C and the supernatant was separated as the soluble fraction. The leftover pellet was resuspended in SDS containing lysis buffer (50mM Tris pH7.5, 150mM NaCl, and 0.1% SDS), sonicated, and centrifuged at 14,000g for 15mins at 4°C. The supernatant was separated and designated as the insoluble fraction. Both the soluble and insoluble fractions were further loaded on an SDS-PAGE and were either stained by silver staining or transferred onto a nitrocellulose membrane and probed with an anti-Orb2 antibody.

### Silver Staining

Soluble and insoluble fractions were run on a 10% SDS-PAGE. The gel was then washed with distilled H_2_O for 5 mins followed by putting it in a fixing solution (50% methanol, 12% acetic acid, 0.05% formaldehyde) for 1 hour. The gel was then washed 3 times with 50% ethanol for 20 mins followed by sensitization with 0.8mM Sodium Thiosulfate for 2 minutes, washing with distilled H_2_O and putting in staining solution (20mg/ml AgNO_3_, 0.076% formaldehyde) for 15 mins in dark. The stained gel was then washed briefly with distilled H_2_O and put in a developing solution (6% w/v Na_2_CO_3_, 2% of 0.8mM sodium thiosulfate, 0.01% formaldehyde) till bands are seen. Then gel was given a wash with distilled H_2_O and put in the stop solution (50% methanol, 12% acetic acid). The gel was then scanned to record the data. Densitometric intensities of individual wells were analyzed with ImageJ software and graphs were plotted using GraphPad Prism software.

### Immunoprecipitation from flies

For immunoprecipitation from fly heads, 100–150 adult fly heads were homogenized in lysis buffer containing 150 mM NaCl, 50mM Tris, 1% NP40, and protease inhibitors. Lysates clarified by centrifugation at 10,000g for 15 minutes at 4°C were incubated with 5ul of serum containing antibodies for 2 hours at 4°C followed by the addition of pre-blocked (with 1% BSA) Protein-A beads (Repligen) for an additional 2 hours. The beads were next washed three times for 10 min with cold lysis buffer and bound proteins were eluted by the addition of 0.2 M Glycine (pH 2) for 1 min and further neutralization with 1.5 M Tris (pH 9). These samples were further subjected to SDD-AGE and western blot.

### SDD-AGE protocol

Samples were mixed with 4x SDD-AGE loading dye and were run at 50V in a 1.3% agarose gel in 1XTAE buffer containing 0.1% SDS. The gel was transferred overnight to a nitrocellulose membrane using capillary transfer with 1X TBS buffer. The membrane was further processed for western blot using anti-Orb2 antibody.

### Immunostaining and imaging

Transfected S2 cells were plated on coverslip bottom dishes and GFP and RFP coexpressing cells were directly imaged on a confocal microscope. For immunostaining experiments, these cells were fixed with 4% PFA for 5 minutes, washed 3X with PBST buffer, followed by blocking for one hour in blocking buffer (5% Normal goat serum (NGS) in PBST buffer). Primary antibodies were prepared in the blocking solution which was added to the cells and incubated for 2 hours at room temperature. Post removal of the antibody solution, cells were washed again 3X with PBST buffer and incubated with the secondary antibody tagged with fluorophores in block solution for 1 hour at room temperature. Next, these cells were washed 3X with PBST and then imaged on a confocal microscope. For immunostaining of the Drosophila brains, a similar protocol was followed, except, the fixation was done for 2 hours and the primary and secondary antibody incubations were done overnight, and the brains were mounted with Vectashield between glass coverslip and slide. For the muscle staining, third instar larvae were dissected, fixed for one hour, washed 3X with PBST, and then stained overnight with fluorophore-tagged phalloidin.

### Fly stocks, handling, and maintenance

Flies were raised on standard cornmeal food in a 12:12 hours light, dark diurnal cycle at 22° C. The following lines were used in this work.

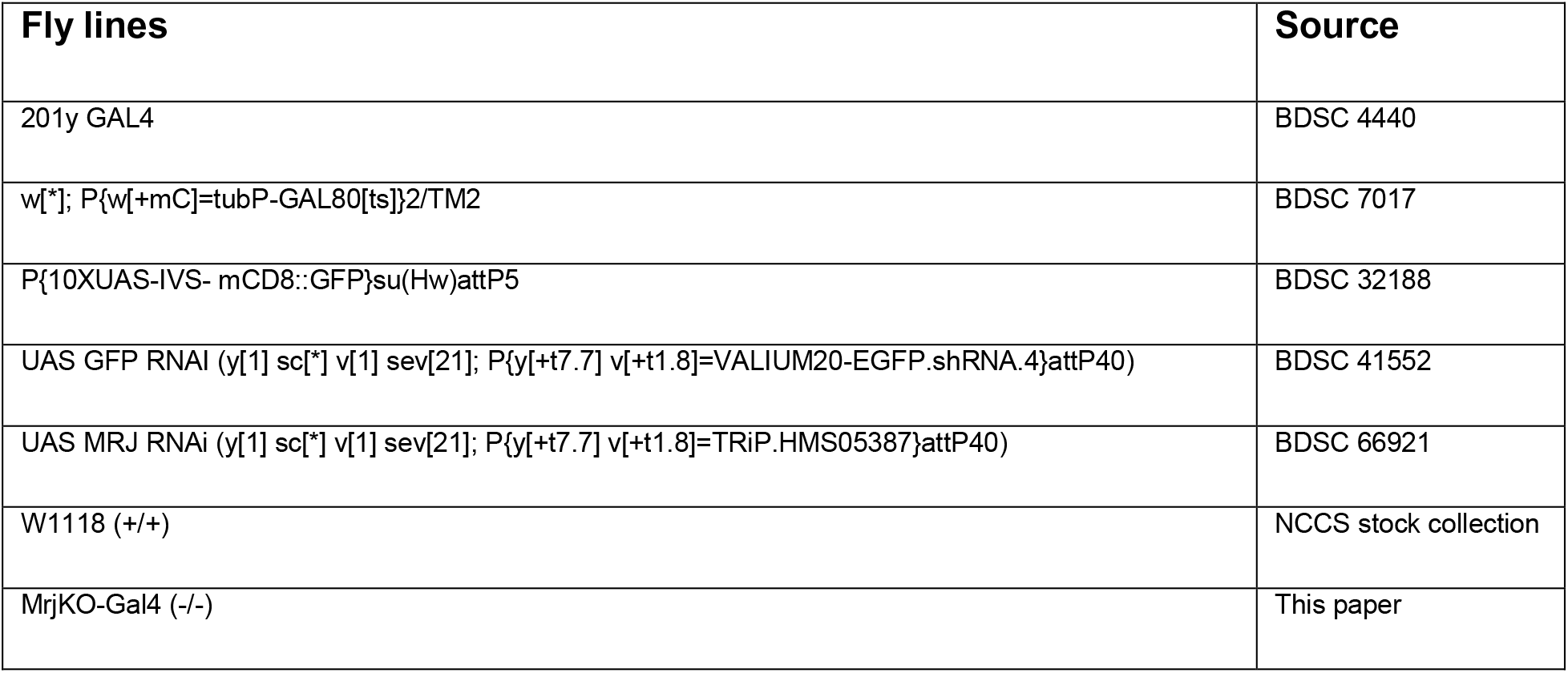

### Adult Fly locomotion assay

Single flies were put inside 8cm long, 0.5cm diameter transparent tubes using an aspirator, and both sides were sealed with cotton plugs. The tubes were tapped against the table to get the fly to one side, followed by putting the tubes horizontally, next to a scale, and recording a video of the fly’s movement. The video was analyzed to measure the speed by measuring the distance traveled to the time.

### Male courtship suppression-based memory assay

Male courtship suppression assays were done as described earlier (Majumdar et al., 2012). Briefly, virgin male flies were made to undergo 3x training with mated females. Each training was of 2 hours duration and there was a gap of 30 minutes where the trainer female was removed. Virgin males which did not undergo any training were used as naïve males for control. Post-training, the male flies were again transferred to new tubes and after different intervals of time, were put into a courtship chamber containing a mated female. Videos of these encounters were recorded for 10 minutes and further manually analyzed to calculate the courtship index as per the following formula:

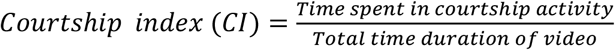

The memory indexes for individual flies were further calculated using:

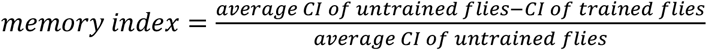

The average memory index was calculated for each genotype at different time points and further plotted using Graphpad prism7.

### Polysome analysis

Polysome analysis was performed on S2 cell extract as per previously published methodology (Joag et al., 2020). Briefly, cycloheximide-induced translationally stalled cells were lysed in polysome lysis buffer (300mM NaCl, 50mM Tris.Cl (pH:8), 10mM MgCl_2_, 1mM EGTA, 1% Triton-X100, 0.02% sodium deoxycholate). The lysate was centrifuged at 10000g for 10 minutes at 4°C, and the resulting supernatant was loaded on a 5-45% sucrose density gradient made in resolving buffer (140 mM NaCl, 25mM Tris-Cl pH:8, 10mM MgCl_2_). This was further spun at 4°C at 28000 g speed for 2 hours on a swing bucket SW41 rotor. The gradient was next fractionated with monitoring of Absorbance at 254 nm using a Biocomp fractionator station. Proteins from all the fractions were precipitated by overnight incubation with 10% trichloroacetic acid at −20° C. Precipitated proteins were spun down at 12500g for 15 min and then washed with chilled acetone. Washed protein pellets were resuspended in 2X loading buffer and subjected to western blot.

### Antibodies and other reagents

The following commercially available antibodies were used in this paper: Anti-HA ab9110 (Abcam), Anti-HA ab18181 (Abcam), Anti-Flag F1804 (Sigma), Anti-Huntingtin Mab2166 (Chemicon), Anti-GFP 50430-2-AP (Proteintech), Anti-RFP R10367 (Life Technologies), Anti α-Tubulin 66031-1 (Proteintech), Anti-FAS II-1D4 (DSHB), Anti-Puromycin-3RH11(Kerafast), Anti-Ref2P-ab178440 (Abcam), Anti-HA epitope tag clone 16B12 (Biolegend), Multi Ubiquitin chain monoclonal-cloneFK2 14220 (Cayman), Alexa flour 488 Phalloidin-A12379 (Life Technologies), Alexa flour 555 phalloidin-A34055 (Life Technologies), Alexa fluor 555 anti-mouse A21424 (Life Technologies), Alexa fluor 488 anti-mouse A11029 (Life Technologies), Alexa fluor 555 anti-Rabbit A11034 (Life Technologies), Alexa fluor 488 anti-Rabbit A21429 (Life Technologies).

Anti-Orb2 and anti-Mrj antibodies were raised against recombinant 6X Histidine-tagged full-length protein, in Rabbit and Guinea pig for Orb2 and Guinea pig for Mrj. For immunoprecipitation, protein-A Agarose beads (Repligen 102500-03) and RFP trap magnetic beads (Chromo Tech rtma) were used.

## Author contributions

Meghal Desai made the chaperone expression library and performed the IP-based screen, and biochemical characterization of Mrj and Mrj knockout and polysome experiments. Hemant performed the immunostainings, confocal imaging, life-span, mobility, and Drosophila courtship suppression-based memory assay. Jagyanseni Naik purified Mrj and performed the DLS experiment. Ankita Deo performed the Yeast Sup35 based prion assays. TB supervised the Yeast experiments along with obtaining the funding for the work. AM conceptualized, and designed the experiments, supervised the work, obtained funding, and drafted the paper with inputs from all other authors.

## Acknowledgment

This work was supported by a Wellcome Trust-DBT India alliance grant (IA/I/13/2/501030), a DBT grant (BT/PR25893/GET/119/174/2017), and NCCS intramural funding to AM. Work in Dr. Tania Bose’s lab was supported by a DBT Ramalingaswami fellowship (BT/RLF/Re-entry/54/2013) and an IYBA grant (BT/09/IYBA/2015/03). We thank Simon Alberti and Randall Halfmann for the yeast strains and the vectors for the chimeric Sup35 prion experiment and the yeast destination vector was a gift from Susan Lindquist’s lab. The Mrj knockout was made in collaboration with Well Genetics. We acknowledge Kausik Si for the Orb2, Konrad Basler for pUASgHA attB, Aravind Penmatsa for the pFastBac-GFP-His, Troy Littleton for the HttQ138, and Norbert Perrimone for the Htt Exon1-GFP constructs used in this paper. Gateway destination plasmids from Drosophila Genomics Resource Center (NIH grant 2P40OD010949), monoclonal antibodies from the Developmental Studies Hybridoma Bank, created by the NICHD of the NIH and maintained at The University of Iowa, and fly stocks from the Bloomington Drosophila Stock Center (NIH P40OD018537) were used in this study. We thank Rishov Goswami, Vishal Naik, Harshada Gade, Maitheli Sarkar, and Vighnesh Ghatpande for being associated during the initial phases of this work and George Fernandes for helping in making the fly food, stock, and cell line maintenance. We also acknowledge Rahul Bankar, animal house, NCCS for his help in generating the antibodies here and Deepa Subramanyam, Vasudevan Seshadri, Gunther Hollopeter, Shreejita Chatterjee, Neelanjana Das, and Gaurav Agarwal for reading and providing valuable suggestions.

---

**Supplementary Figure 1:**
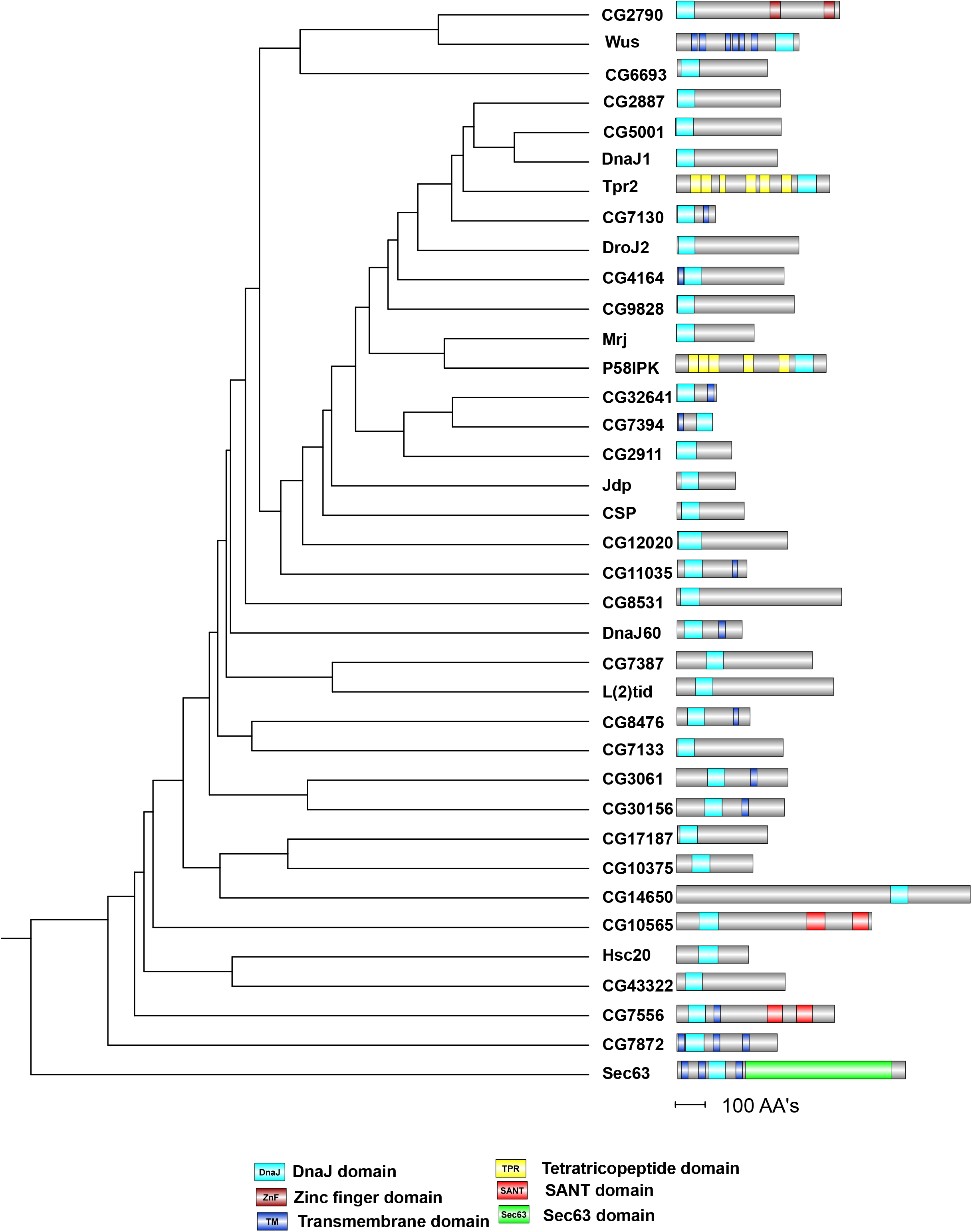
Rooted phylogenetic tree with branch length for all the Hsp40 family of proteins along with their predicted domain structures. This list does not show Auxillin and Rme8, which were not used in the screen here.

**Supplementary Figure 2:**
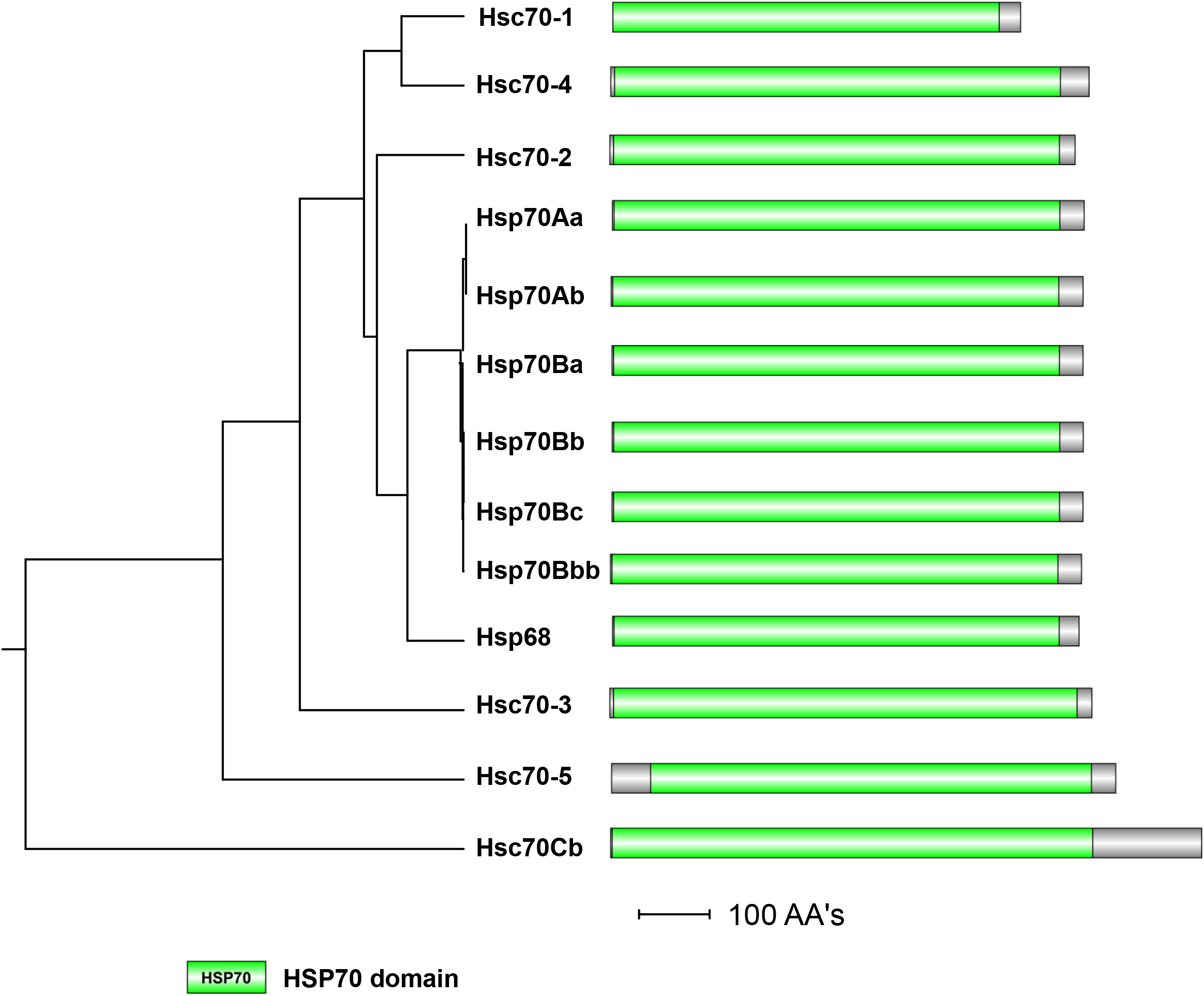
Rooted phylogenetic tree with branch length for all the Hsp70 family of proteins along with their predicted domain structures. Of these, Hsp70Aa, Hsc70-1, Hsc70Cb, and Hsc70-4 were used in the screen.

**Supplementary Figure 3:**
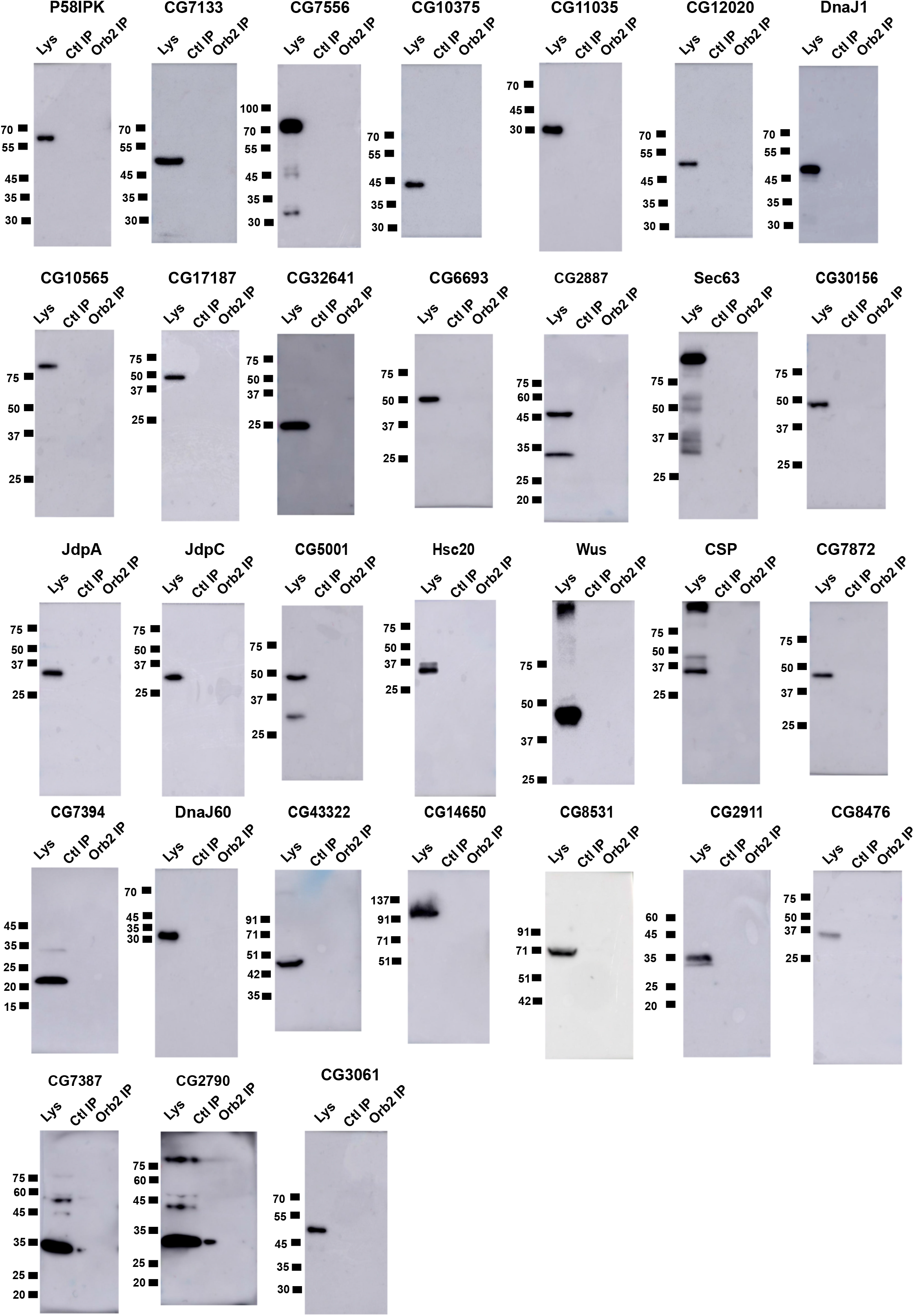
Representative western blots of 31 Hsp40 proteins, which don’t show interaction with Orb2A in the immunoprecipitation screen.

**Supplemental Figure 4:**
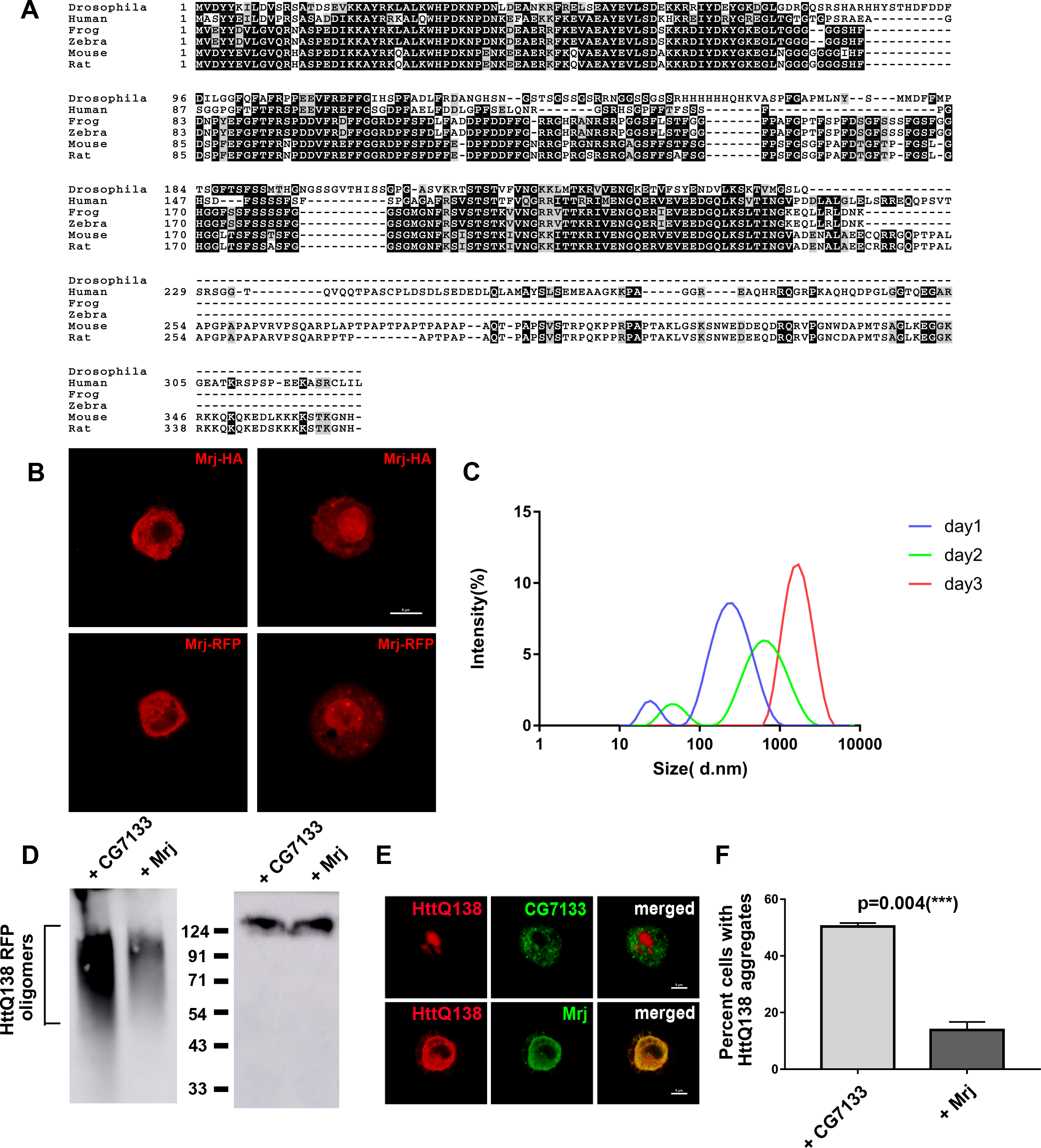
**A**. ClustalW alignment of Drosophila Mrj with Human, Frog, Zebrafish, Mouse, and Rat Mrj/DnaJB6 **B**. Representative images of S2 cells expressing Mrj-HA (upper panels) and Mrj-RFP (lower panels). Both constructs show the presence of Mrj in both nucleus and cytoplasm **C**. DLS experiments with recombinant Mrj over three days show its shift to higher sizes with time **D**. Left panel shows a representative SDD-AGE from S2 cell lysate coexpressing HttQ138-RFP along with CG7133 and Mrj showed a decreased amount of Htt oligomers in presence of Mrj. The right panel is of a western blot of lysates in SDS-PAGE from S2 cells coexpressing HttQ138-RFP with Mrj and CG7133 showing similar amounts of Htt **E**. Representative images of HttQ138-RFP cells coexpressing with CG7133-HA and Mrj-HA suggest a decrease in the Htt aggregates in presence of Mrj **F**. Quantitation of the percentage of HttQ138-RFP expressing cells with aggregates in presence of CG7133 and Mrj suggests a significant decrease of Htt aggregates in presence of Mrj. Data is represented as Mean ± SEM and unpaired t test is used here to check significance.

**Supplemental Figure 5:**
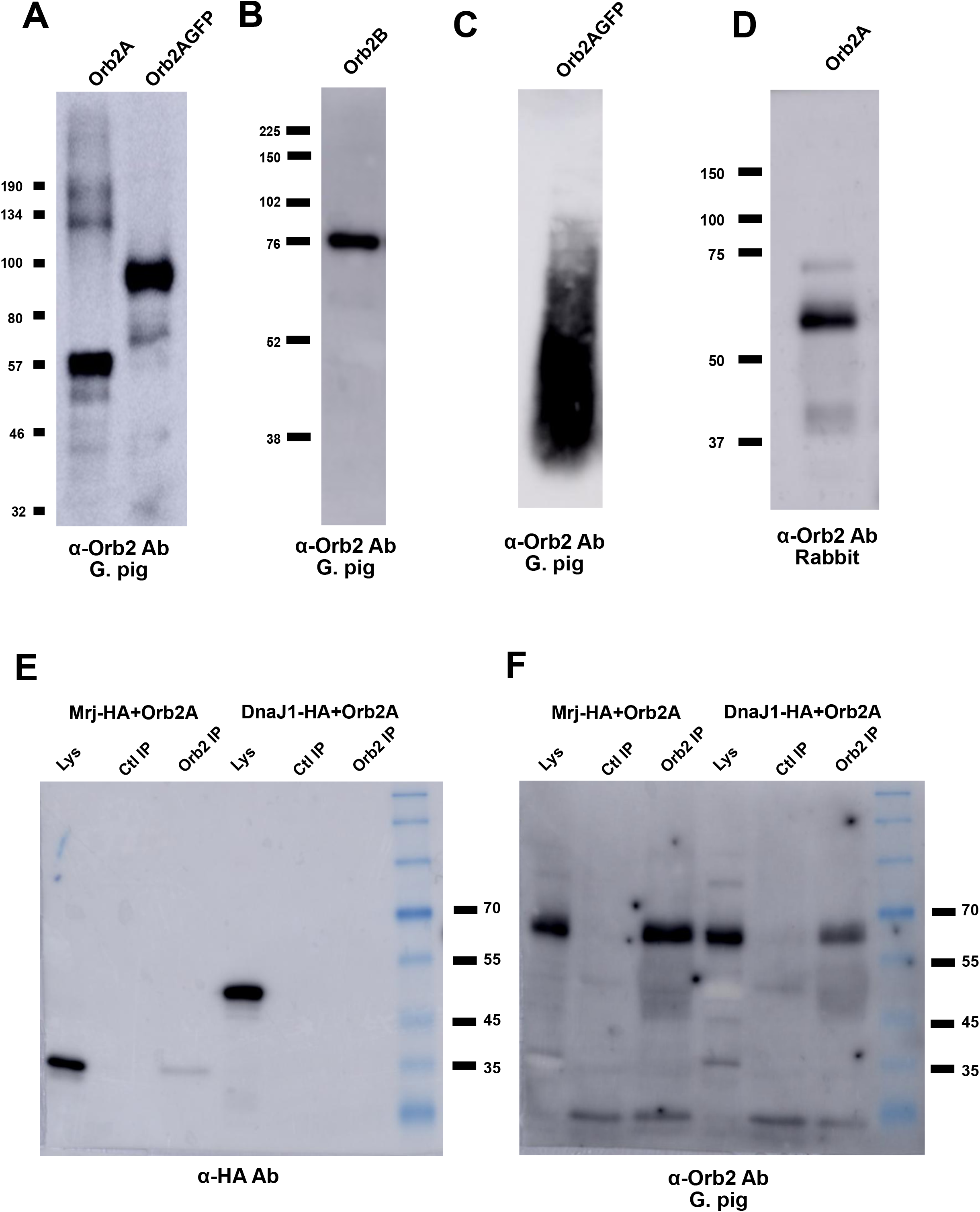
**A**. Validation of anti Orb2 antibody raised in guinea pig in western blot with S2 cell lysate from cells expressing Orb2A and Orb2A-GFP **B**. Guinea pig anti-Orb2 antibody detects endogenous Orb2B from fly head extract **C**. Guinea pig anti-Orb2 antibody detects Orb2A-GFP oligomers from Sf9 cell lysate **D**. Validation of anti Orb2 antibody raised in rabbit in western blot with S2 cell lysate from cells expressing Orb2A **E**. Representative western blot of immunoprecipitation performed from cells expressing Orb2A with Mrj-HA and Orb2A with DnaJ1-HA with anti Orb2 antibody. Probing the blot with an anti-HA antibody shows the presence of Mrj in the lane for lysate and Orb2 IP suggesting its interaction with Orb2A. In contrast for DnaJ1, it is detected only in the lysate lane and not in the Orb2 IP lane, suggesting no interaction between DnaJ1 and Orb2A. **F**. The same blot as in E was probed with anti Orb2 antibody and here Orb2A could be detected in both the lysate lanes and Orb2 IP lanes confirming the pull down of Orb2A with anti Orb2 antibody.

